# The kinase inhibitor Palbociclib is a potent and specific RNA-binding molecule

**DOI:** 10.1101/2022.01.20.477126

**Authors:** Matthew D. Shortridge, Venkata Vidalala, Gabriele Varani

## Abstract

The growing awareness of the role of RNA in human disease has motivated significant efforts to discover drug-like small molecules that target RNA. However, high throughput screening campaigns report very low hit rates and generally identify compounds with weak affinity, while most structures reported in Academic studies also lack the pharmacological properties of successful drugs. Even FDA-approved RNA-targeting drugs have only weak (10 μM) binding activity. Thus, it is often stated that only complex RNA structures, such as the ribosome or riboswitches, are amenable to small molecule chemistry. We report that the kinase inhibitor Palbociclib/Ibrance is a nM ligand for the HIV-1 TAR. It inhibits recruitment of the positive transcription elongation factor complex at nM concentrations and discriminates >20 fold. We further show that RNA binding can be fully decoupled from kinase inhibition, yielding a new molecule with even higher affinity for RNA. We thus demonstrate that nM affinity, specificity, and potent biochemical activity against ‘undruggable’ RNAs can be found in the chemical space of blockbuster drugs.

## Introduction

Targeting RNA with drug-like small molecules would provide new therapeutic avenues to addressing many unmet clinical needs. However, the chemistry and pharmacology of small molecule binding to RNA is under-explored, with the exception of natural product antibiotics which target ribosomal RNA^1–3^. Despite very substantial recent efforts in the biotech and pharma industries^4,5^, and the success of splicing modifiers developed at Roche and Novartis^6–8^, the identification of pharmaceutically attractive small molecules which bind to RNA potently and specifically remains a very significant challenge^2,3,9^. Owing to the difficulty of achieving specificity sufficient to discriminate between RNAs, it has been suggested that only complex RNA structures, such as riboswitches or the ribosome, are amenable to inhibition with drug-like small molecules^2^. Here we show that the blockbuster drug Palbociclib/Ibrance binds to HIV-1 TAR, a simple RNA stem-loop, with potent (100 nM) and specific activity and is active in rigorous biochemical assays at low nM concentration.

Ribosomal RNA and riboswitches provide some of the best known examples of recognition of structured RNAs by small molecules^1,10^. However, the elaborate higher order folds observed in these RNAs are not common in non-coding RNAs and mRNAs, whose structures are frequently destabilized by single strand RNA binding proteins and during active translation^11,12^. Simpler secondary structures such as stem loops and internal loops are ubiquitous in cellular and viral RNAs and perform well-established regulatory functions by regulating pre-mRNA splicing, 3’-end processing, protein synthesis, etc. These secondary structures persist in the cell when more complex structures are less stable^13,14^. However, because they are so superficially similar to each other and devoid of binding pockets, specific targeting of these RNAs with drug-like small molecules is often described as unlikely to succeed^2,15^.

An extensively system for the study of interactions of small molecules with RNA is TAR, where inhibiting its interaction with Tat protein has long been pursued in efforts to discover new anti-viral agents^16^. The TAR UCU bulge region is recognized by the arginine rich motif (ARM) of the HIV protein Tat, to recruit the host super elongation complex (SEC) to the apical loop and enhance proviral transcription (Fig. 1). Although multiple TAR ligands have been reported in the academic literature, most of these molecules have lacked rigorous evidence of cellular or biochemical activity and the pharmaceutical characteristics of known drugs, with a few notable exceptions^17–20^. We have shown that this RNA can be targeted with very high affinity (30 pM) and specificity (>10,000 fold) by macrocyclic peptides^21–23^. However, even these ultra-potent peptides only weakly inhibit recruitment of P-TEFb to TAR and therefore reactivation of transcription^23^.

**Figure 1.**
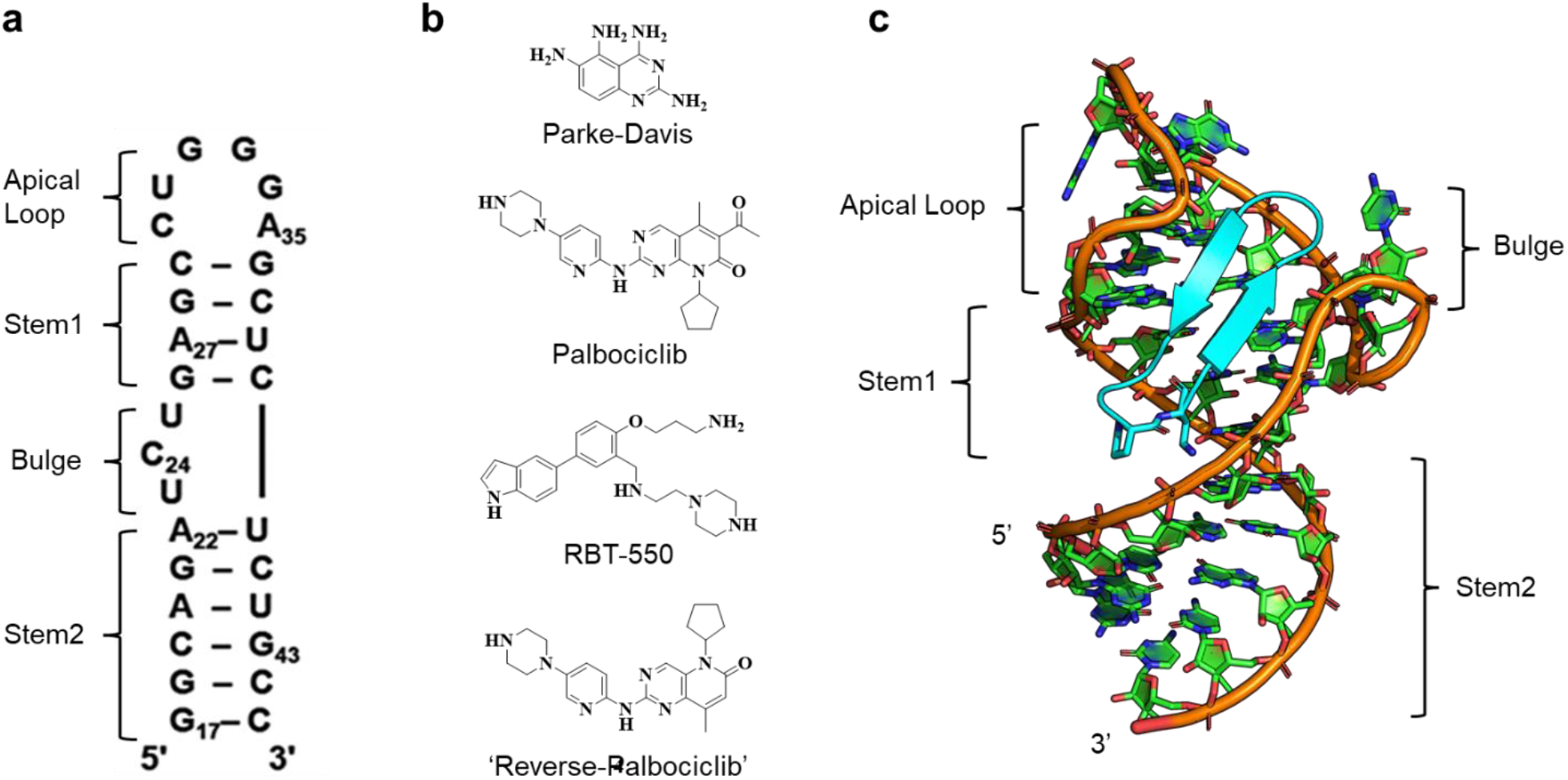
a) Secondary structure of the HIV1-TAR RNA. b) The Parke-Davis group reported compounds which bind to the loop region of HIV-TAR (top). The amino pyrimidine core structure of the Parke-Davis fragment is found in several FDA-approved drugs, including Palbociclib (second from top) and Ribociclib; the third structure from the top is RBT-550 from the Ribotargets project^17^, while the compound at the bottom is ‘Reverse Palbociclib’, as described in the main text. C) The three-dimensional structure of the HIV-TAR RNA in the presence of peptide JB-181, showing that the UCU bulge and apical loop are juxtaposed to each other, creating a new binding pocket that is absent in free TAR^23^.

Here, in contrast, we report that the blockbuster drug Palbociclib/Ibrance binds to HIV TAR with a KD of approximately 100 nM, specifically recognizes the TAR apical loop and UCU bulge simultaneously and disrupts formation of the SEC-TAR complex at low nM concentration. Remarkably, the affinities of Palbociclib for TAR and for cdk6 are comparable, even though the latter result was achieved only after a decade-long medicinal chemistry effort^24^. In addition, we show that RNA and kinase binding can be decoupled, demonstrating that binding of Palbociclib to RNA is in fact unrelated to kinase inhibition. Thus, we demonstrate that targeting of ‘undruggable’^2^ RNA secondary structures can be accomplished with pharmaceutically attractive small molecules.

## Results

The concept that drug-like small molecules can bind to RNA potently and specifically is often met with skepticism by experienced medicinal chemists and indeed several big pharma projects in this area have been terminated. Many academic manuscripts have described small molecules that bind to RNAs, but very seldom have affinities stronger than 10-100 μM been reported, unless the chemical space of known drugs is abandoned through multivalency^25^ and/or high basic charge^1^, with a few notable exceptions^26,27^. Even for HIV TAR, a paradigmatic system studied for more than 2 decades, molecules reported to bind to it with sub-μM affinity (aminoglycosides and other basic molecules, flexible short peptide mimetics, substituted acridines, etc.^28,29^) have almost invariably lacked the properties of successful drugs. A few exceptions stand out: namely, the small molecule RBT550 and related series^17,18^, and several fragments reported by Parke-Davis^19,20^. The 2,4,5,6-quinazolinetetramine core structure^20,30^ (Fig. 1b) identified in fragment-based searches, targets the apical loop of TAR; this is important, because small molecules that bind there would inhibit P-TEFb, while molecules which bind to the UCU bulge where Tat protein binds would be unlikely to elicit a biological response^23^. Furthermore, a ligand that binds to the flexible loop could generate its own binding pocket by refolding the RNA around it, as has been observed in the HepC IRES^26,27^ and with our pM peptides^23^ (Fig. 1c).

Despite this premise, it is highly unlikely that fragments such as the Parke Davis compounds would bind RNA potently and specifically, as they are too small. Thus, we searched for larger chemical structures which contain the features of these fragments. Two such molecules are the FDA-approved breast cancer drugs Palbociclib/Ibrance (Fig. 1b; PD0332991, a Parke-Davis project^24^), and Ribociclib/Kiskali (Supplementary Fig. 1), both cdk4/cdk6 inhibitors. We show herein that Palbociclib binds to HIV TAR with 100 nM affinity, comparable to its affinity for cdk6^24^, a remarkable result, and that it discriminates >20 fold against another stem-loop, pre-miR-21.

### Palbociclib binds to HIV TAR RNA with nM affinity

To detect binding to RNA, we used an unbiased NMR approach based on relaxation editing because it is universally applicable to any biomolecule without the need to set up a target-specific assay^31^. We have shown that this method works well with RNA and reasoned that this method would easily detect μM binding, which we expected to find^32,33^. Compounds which bind are identified through changes in peak height when comparing spectra for RNA-free and RNA-bound ligands. The decrease in peak height is proportional to the increase in ligand linewidth relative to the free small molecule, which occurs because the bound small molecule rotationally diffuses like the much larger biomolecule it is bound to, leading to a large decrease in the relaxation rate T2. This method depends only on the much longer rotational correlation time of the bound vs free small molecule and is much less prone to false positives than other NMR methods used to screen for small molecules binding to RNA^34–36^.

We applied this approach to the cdk4/cdk6 inhibitors Palbociclib, Ribociclib and Abemaciclib, among other compounds, examining them for binding to HIV TAR and to pre-miR-21, two well-known RNAs of therapeutic interest. These two RNAs are similar in secondary structure, shape and size, but have divergent sequence and local structure and were therefore chosen to assess selectivity. At 10 μM concentration, both RNAs showed a clear response in binding Palbociclib, as demonstrated by the fact that the ligand signals are essentially broadened into the noise (100 μM small molecule and 10 μM RNA in bis-Tris buffer at pH 6.5; Supplementary Fig. 2). Under these conditions, the other two inhibitors, Ribociclib and Abemaciclib, also bind to both RNAs (data not shown).

To provide a quantitative estimate of binding, we titrated 100 μM samples of Palbociclib with either HIV TAR or pre-miR-21 over the same RNA concentration range (up to 5 μM) (Fig. 2). In order to assess whether binding is driven by electrostatic interactions mediated by the charge on the piperazine ring (predicted pKa of 8.5), we used a buffer containing 50 mM d19 bis-Tris at pH 6.5, with the addition of 200 mM NaCl, 50 mM KCl and 4 mM MgCl2 to mimic the ionic environment of the cell. NMR data were collected and processed as described in methods, with spectra normalized to the non-binding internal reference DSA. For both HIV-TAR (Fig. 2c) and pre-miR21 (Fig. 2d), the free small molecule spectrum is shown at the bottom, with spectra recorded at increasing RNA concentrations (0.1-5 μM) stacked above. The rapid decrease in the Palbociclib signal upon the addition of HIV TAR demonstrates high affinity, on the order of a few hundred nM, whereas changes in ligand signal are only observed at the highest concentrations of pre-miR-21, indicating an affinity of the order of 10 μM, a >20-fold difference.

**Fig. 2.**
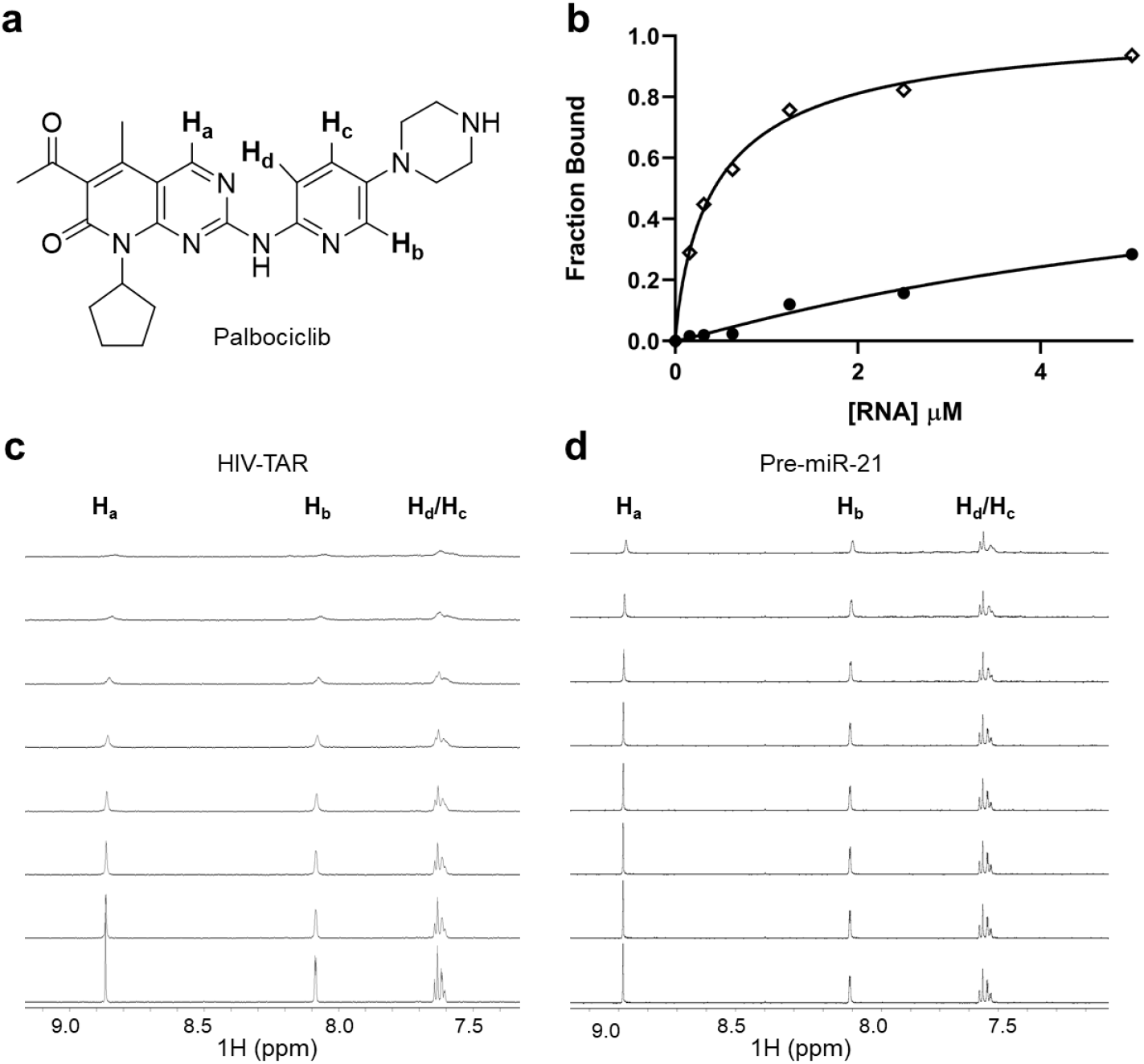
NMR titration of Palbociclib with TAR RNA and pre-miR-21. a) Structure of Palbociclib; b) Palbociclib peak intensities are plotted as a function of RNA concentration, with fits obtained using equation 5 (see methods and reference 31) Spectra were normalized to the non-binding internal reference standard sodium 4,4-dimethyl-4-silapentane-1-sulfonate (DSA; 9 protons) and intensities were used to generate a binding isotherm, from which approximate binding constants were generated by curve fitting (◊ HIV TAR, • Pre-miR-21). c, d) 1H NMR titration series. 100 μM Palbociclib was titrated with HIV TAR in 10 μL steps of 500 μM RNA stock solution (0-5 μM) (c) and with pre-miR-21 RNA (0-5 μM) (d). These spectra were collected at 800Mhz and 37 °C, in 50 mM d9 deuterated bis-tris buffer at pH 6.5, containing 200 mM NaCl, 50 mM KCl and 4 mM MgCl2.

### Palbociclib binds to RNA potently in both fluorescence and NMR assays

Binding affinities can be quantified by applying equation 5 (see experimental section) to fit the signal intensity as a function of RNA concentration^31^ (Fig. 2b). Using this approach, we calculate an approximate affinity for Palbociclib binding to HIV TAR of 335±50 nM with a Bmax greater than 0.9, suggesting a fully saturated and specific 1-to-1 complex, consistent with NMR titrations (see below). Palbociclib binds instead to pre-miR-21 with an affinity of 7,000±5,000 nM. The very large uncertainty, coupled with the small decreases in ligand signal and failure to reach saturation, suggests that Palbociclib binds to pre-miR21 non-specifically. This conclusion is supported by the observation that the difference in binding affinity toward the two RNAs is reduced in low salt buffer (100 nM for TAR vs 1 mM for pre-miR-21); in contrast, high affinity for TAR is retained when high ionic strength conditions are introduced to decrease electrostatic binding, while binding to pre-miR-21 is more strongly affected by ionic conditions.

We used a fluorescence assay to more rigorously quantify binding of Palbociclib to HIV TAR, with U25 substituted by 2-aminopurine (2-AP)^37^. The binding isotherm shows a rapid increase in fluorescence, consistent with an increase in 2-AP base stacking, followed by a slower decrease in intensity when the ligand concentration exceeds 200 nM, suggesting the presence of an additional binding mode (Fig. 3 Top). Similar increases in fluorescence are observed with Neomycin, Tat^37^ and with cyclic peptides with pM-nM affinity (Supplementary figure 2 in reference 23); this same biphasic behavior has also been reported for Neomycin and Tat peptides^37^. The high affinity binding of Palbociclib to HIV TAR corresponds to a K_D_=104±80.2 nM when the curves were fit to a two-site binding mode (equation 1 in the methods section), a value very close to that obtained by NMR relaxation titrations under similar ionic conditions. This result was obtained in the presence of a large excess of competitor tRNA, 250-fold, which we routinely use to reduce non-specific binding. The secondary low affinity binding site was found to have an affinity greater than approximately 700 nM.

**Fig. 3.**
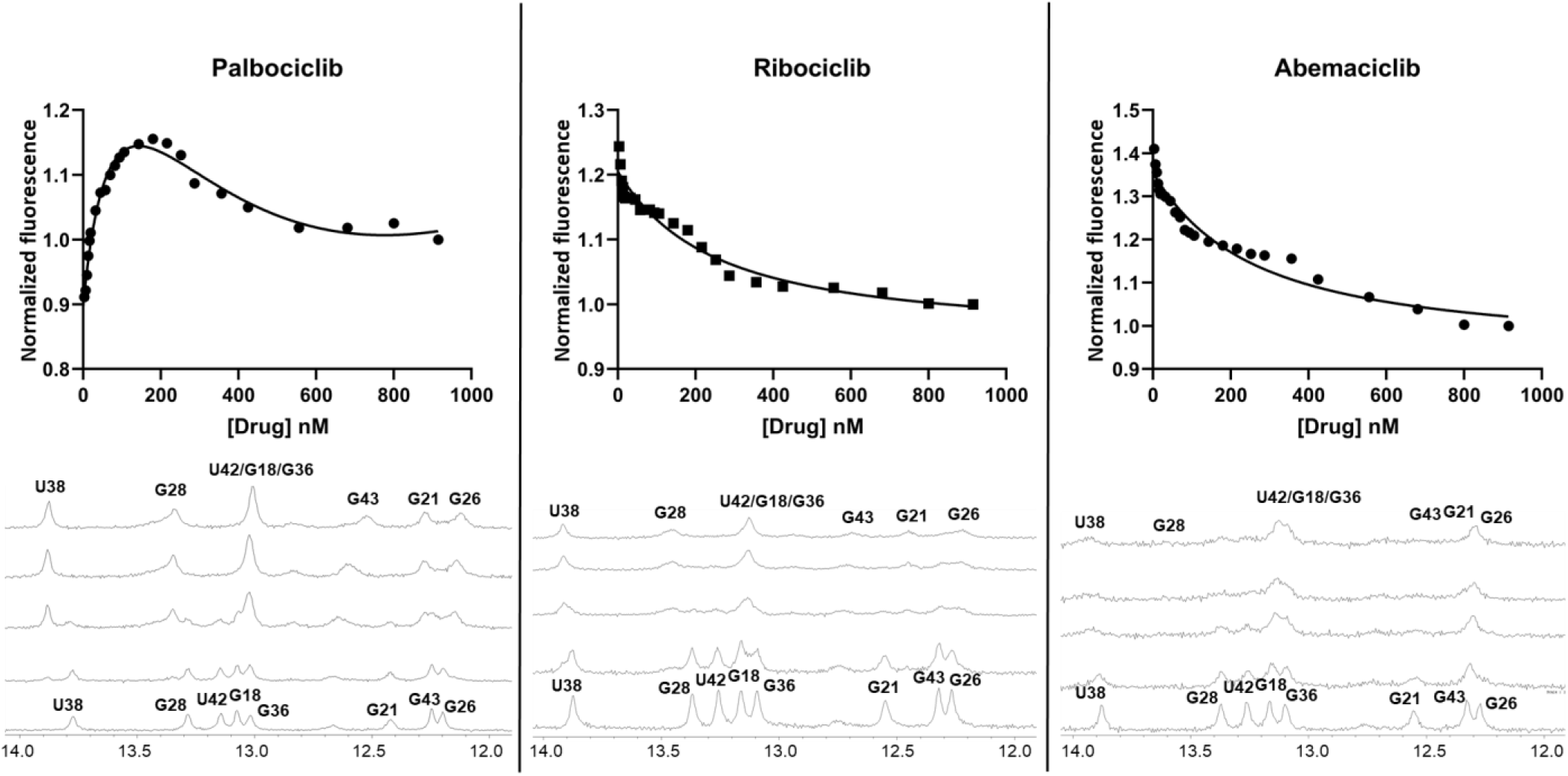
Binding of Palbociclib and two other cdk4/cdk6 kinase inhibitors to HIV TAR, monitored by fluorescence and NMR. (Top) 2-amino-purine (2-AP) HIV TAR binding assays and (Bottom) 1D imino-region NMR titrations for Palbociclib, Ribociclib and Abemaciclib. For Palbociclib, the 2-AP assay shows a rapid initial increase in fluorescence signal, indicating an increase in base stacking for 2-AP upon binding, as has been observed for other ligands^23,37^. This increase is followed by a decrease in the fluorescence signal at higher concentrations, as previously observed for aminoglycosides and Tat protein^37^. This biphasic binding curve suggests the presence of two sites, a high affinity site with apparent KD of 104±80 nM (under low ionic conditions; 50 mM bis-Tris buffer) and a low affinity site with an apparent Kd >700 nM. In the NMR titrations, Palbociclib binding occurs in the slow exchange regime, consistent with nM affinity, while binding to Ribociclib is in the intermediate-slow regime, with behavior similar to Palbociclib but also significant line broadening. Abemaciclib binds in the intermediate regime, leading to exchange-broadened spectra, consistent with low μM/high nM affinity. The Palbociclib and Ribociclib titrations saturate at a 1:1 RNA-small molecule ratio; the top spectrum in each panel was recorded in the presence of excess small molecule and no further changes were observed.

Ribociclib and Abemaciclib, which bind to the same kinases as Palbociclib, did not display the same response in the fluorescence assays (Fig. 3); instead, only a decrease in the fluorescence signal was observed, suggesting that U25 remains unstacked during binding, similar to what is observed for the weaker binding phases of Neomycin, Tat and Palbociclib. The larger than expected error in these measurements could result from slight differences in tRNA concentrations between measurements, since tRNA would be expected to compete successfully with non-specific binding events.

### Palbociclib binds TAR at a specific binding site, forming a long-lived complex

The fluorescence and NMR relaxation editing results are further supported by the analysis of ^1^H NMR titrations (Fig. 3 bottom). Signals in the RNA imino region saturate near a Palbociclib:TAR ratio of 1:1 and are very clearly in the slow-exchange regime expected for nM binding. For example, the U38 NH resonance, which is well resolved, gives rise to two signals (free and bound) in slow exchange, separated by about 0.1 ppm, corresponding to an off rate much slower than ~60 Hz and therefore a dissociation constant well below μM; the same behavior is observed for several other resonances in the spectrum. The peak shape is comparable to that observed for a 30 nM peptide (Supplementary Fig. 9 in reference 23). In contrast, titration with Abemaciclib results in line-broadened spectra, consistent with intermediate exchange and an off rate approximately equal to the difference in frequency, i.e. the high nM/low μM affinity regime, consistent with the slowly decreasing phase of the fluorescence titration. The NMR response to titration with Ribociclib is intermediate between those of Palbociclib and Abemaciclib, with spectral changes similar to Palbociclib but with increased spectral broadening, as expected because Ribociclib is more closely related to Palbociclib than to Abemaciclib. These results are fully consistent with the affinity estimated from fluorescence and ligand-detected NMR titrations.

In Figure 4, we show TOCSY spectra for HIV TAR and pre-miR-21 and compare them with the spectra of the same RNAs when fully titrated with Palbociclib. The large chemical shift changes for HIV TAR and sharp spectral features confirm that the compound binds in the slow exchange regime and induces large structural changes in the RNA; the chemical shifts are dispersed throughout the molecule, as we observed with the pM cyclic peptides. These large chemical shift changes are not observed for other small molecules that bind to TAR in the ~10 μM affinity range^32^, or for pre-miR-21, where only small chemical shift changes are observed instead (Fig. 4B), together with irregular line shapes indicative of a dynamic, poorly defined interaction site. This conclusion is reinforced by the paucity of intermolecular NOEs in the spectra of pre-miR-21 bound to Palbociclib (Supplementary Fig. 3), which is in contrast with the many (about 100) NOEs observed for the TAR complex. In addition, while the intermolecular NOEs for HIV TAR complexed with Palbociclib are sharp and well-defined, the few NOEs with pre-miR-21 are broad and weak, as expected for a weak non-specific interaction, driven perhaps by the basic charge on the piperazine.

**Fig. 4.**
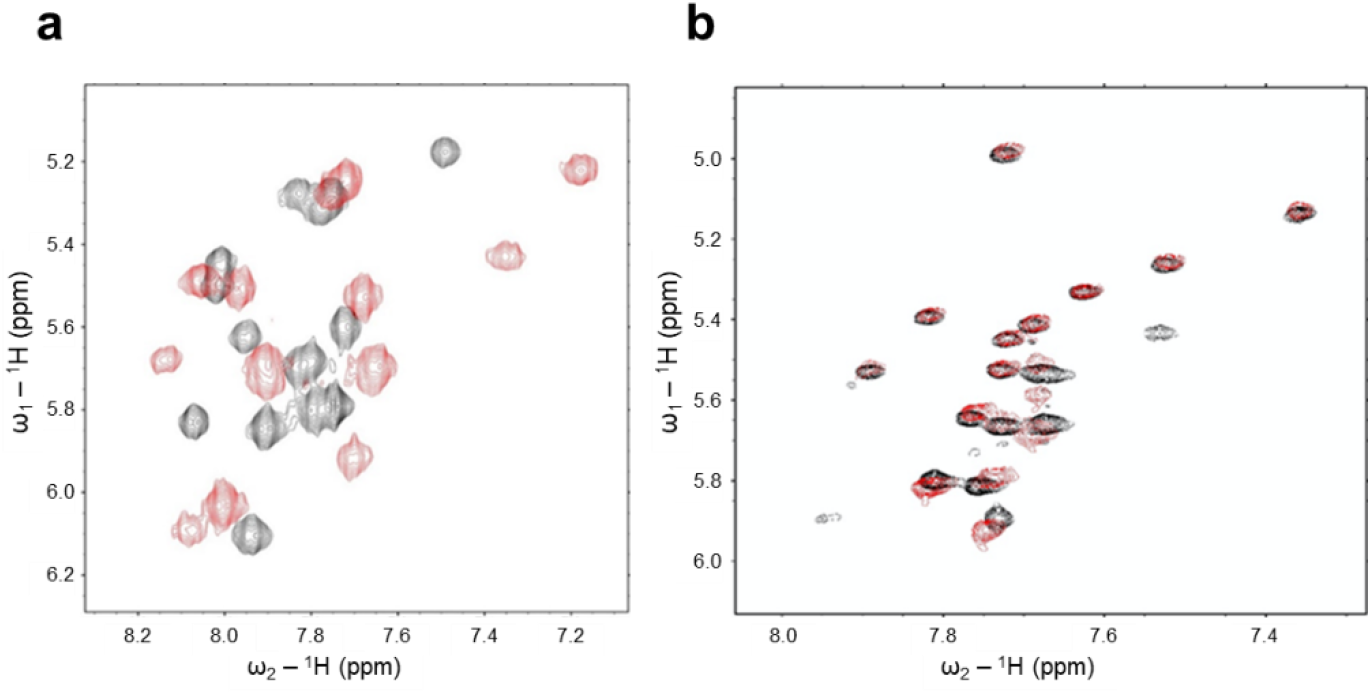
TOCSY spectra of Palbociclib bound to TAR and pre-miR-21. Binding of Palbociclib to the two RNAs occurs with very different affinity, as reflected in the NMR spectra. TOCSY spectra were collected at 800MHz under high salt buffer conditions (200 mM NaCl, 50 mM KCl and 4 mM MgCl2) at 37C. 250 μM RNA (black) was titrated with Palbociclib (red) until saturation was reached (as established from the absence of further changes in the spectra) and changes in chemical shifts were recorded. a) HIV TAR shows dramatic chemical shift changes and ‘slow exchange’ behavior between bound and free forms, with regular peak shapes for most signals, indicative of high affinity and specific binding, whereas b) pre-miR-21 shows much smaller chemical shift changes and irregular peak shapes, indicative of weaker affinity and a dynamic, poorly defined interaction site. These data support the conclusion that Palbociclib binds potently to HIV TAR but much more weakly to pre-miR-21, i.e. that the interaction is specific.

Altogether, we conclude that multiple NMR experiments and a fluorescence titration assay demonstrate that Palbociclib binds to HIV TAR RNA with an affinity of about 300 nM under ionic conditions comparable to the cellular environment, while the functionally related Ribociclib and Abemaciclib bind with much weaker affinity (or a diffuse binding mode). Remarkably, the affinity of Palbociclib for TAR is comparable to what is observed for its original cellular targets, cdk4/cdk6, for which this ligand was optimized over a decade-long effort^24^. Moreover, the high-affinity binding we observe for Palbociclib to HIV TAR remains strong even over relatively large changes in pH (Supplementary Fig. 4)^24^. The discovery of ligands with such high affinity for RNA is rare and has required years-long, sustained synthetic efforts^17,18^.

### Palbociclib inhibits recruitment of the P-TEFb complex

In Supplementary Fig. 5, we overlay TOCSY spectra of free HIV TAR (black) and its complexes with the peptide JB181^23^ (K_d_=30 pm, cyan) and with Palbociclib (red). The Palbociclib chemical shift changes differ considerably from those seen for the JB181 complex, indicating that HIV TAR adopts a distinct structure when bound to it, or at minimum a new binding mode. Notably, the sites of the largest chemical shift changes between free- and Palbociclib-bound TAR map to the apical loop of HIV TAR, rather than the UCU bulge, although both sites are clearly involved (Supplementary Fig. 5). This statement is supported by the preliminary analysis of the NOESY data, which also reveal that binding occurs in the TAR major groove.

The apical loop is the P-TEFb binding site^38^, suggesting that Palbociclib could inhibit formation of the SEC complex containing PTEFb/AFF4/Tat/TAR that activates transcription of the integrated viral genome. To test whether the compound would affect assembly of this complex on TAR, we ran binding shift assays to compare the affinity of the SEC complex to TAR in the presence and absence of Palbociclib (Fig. 5). At 10 nM concentration, we observed a 100-fold decrease in the apparent affinity of the minimal SEC for TAR (K_D_ 36 nM compared to 0.3 nM), even in the presence of 250-fold excess of yeast tRNA. This is a remarkable result; in previous work, we showed that cyclic peptides with 30 pM affinity for TAR had no significant effect on SEC binding (supplementary figure 6 in 23), because they structurally mimic Tat protein. The decrease in affinity of the SEC toward the TAR-Palbociclib complex is comparable to what was observed when the UGGG loop residues of TAR were mutated to CAAA (KD increased from 0.23nM to >10nM)^38^.

**Fig. 5.**
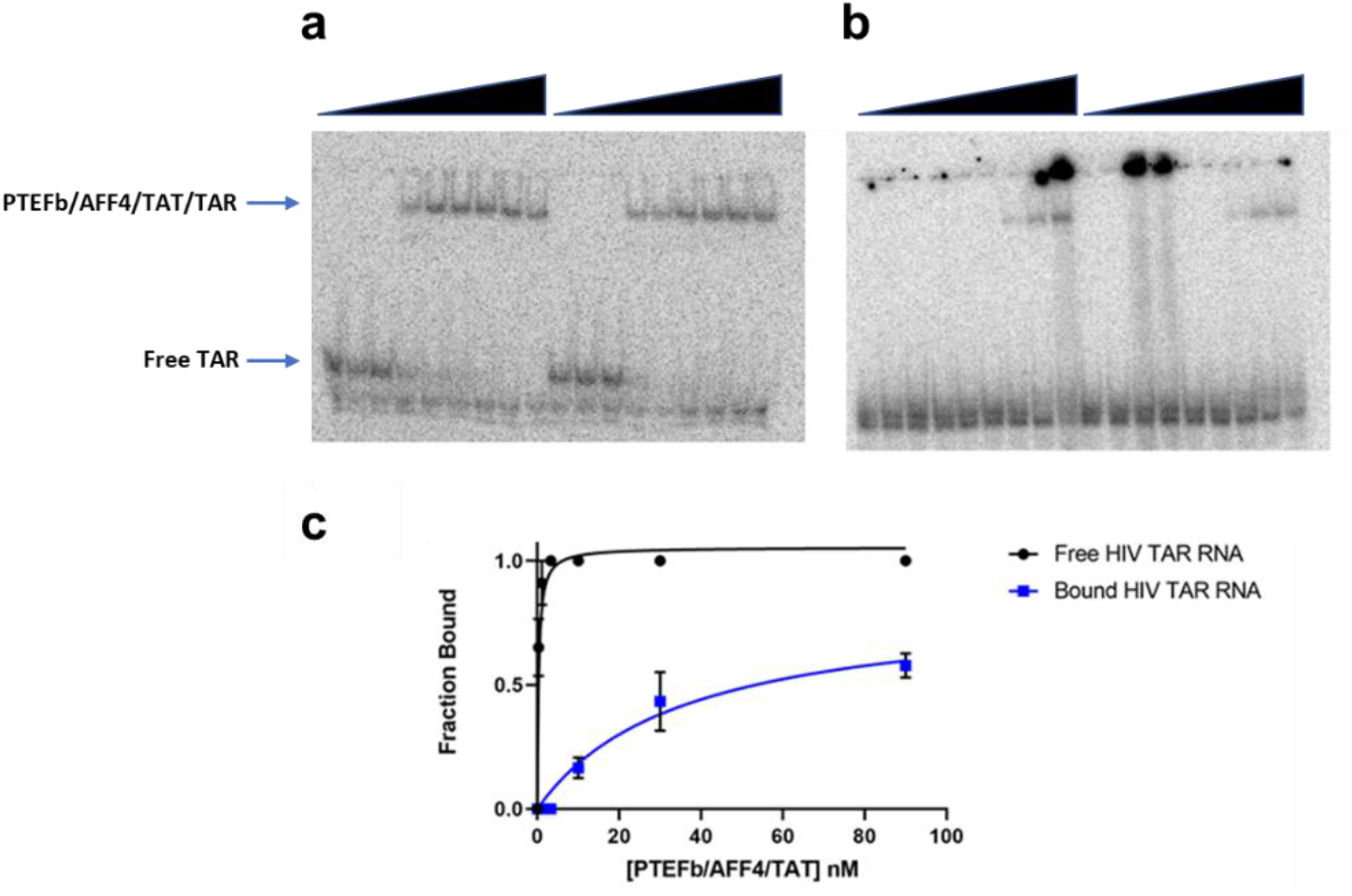
Palbociclib disrupts assembly of the SEC-TAR complex by reducing the affinity between HIV TAR and the core Super Elongation Complex (PTEFb/AFF4/Tat). a) 0.50 nM of 5’ end ^32^P labeled HIV TAR were incubated with 0-90 nM of the preformed PTEFb/AFF4/Tat complex (by serial dilution). b) Palbociclib (10 nM) was pre-incubated with TAR RNA prior to binding (0-90 nM of the PTEFb/AFF4/Tat complex by serial dilution). c) Both assays were repeated in duplicate, as shown, and resolved on 6% native acrylamide gels; bands were quantified with imageJ and plotted vs concentration of the SEC complex. The binding affinity (KD) was calculated by curve fitting to be 0.34 ±0.1 nM for free TAR RNA, but it increased to 35.6 ±10 nM when 10.0 nM Palbociclib were preincubated with the RNA prior to complex formation.

### Binding to RNA is independent of and unrelated to kinase inhibition

It is sometimes stated that kinase inhibitors make good RNA-binding molecules, although the published evidence for this is minimal, but it is unlikely that kinase binding affinity is the reason Palbociclib binds so strongly to HIV TAR. After all, Palbociclib and Abemaciclib (Supplementary Fig. 1), both cdk4/cdk6 inhibitors, bind to RNA with different affinities and characteristics, as revealed by fluorescence measurements and ^1^H NMR titrations (Fig. 3). Only Palbociclib can access the high-affinity (100 nM) site.

To investigate whether a correlation between kinase binding affinity and RNA binding affinity exists for these molecules, we synthesized a compound we call ‘Reverse Palbociclib’, in which the pyrido[2,3-d]Pyrimidin-7-one core fragment of Palbociclib was reverse-linked to the basic piperazine to generate a pyrido[3,2-d]pyrimidin-6-one core coupled to the piperazine in the same manner (Fig. 1b and Supplementary Figure 6; the acetate group was removed to improve synthetic accessibility). Based on modeling studies, we hypothesized that this change in core structure would essentially eliminate binding to the kinases, because the 5-(piperazin-1-yl)pyridin-2-amine would introduce steric clashes within the kinase active site (Supplementary Fig. 6). This hypothesis was tested by assaying inhibition of a panel of 330 human kinase enzymes in the kinome-wide Nanosyn Caliper assay^39^. The results demonstrated only weak inhibition (at 10 μM concentration) of a small number of kinases (<10) by ‘Reverse Palbociclib’ (Supplementary Table 1).

Having excluded kinase binding for ‘Reverse Palbociclib’, we investigated its RNA-binding activity using the ligand-detected NMR relaxation assay (Supplementary Fig. 7) and measured 150±10 nM affinity for the new compound with TAR in high salt buffer, and approximately 10 nM in bis-Tris buffer at pH 6.5 without added counterions. This new ‘Reverse Palbociclib’ structure also improved binding affinity to pre-miR-21 by about 20-fold, showing an apparent affinity of 0.36 ± 0.1 μM, while retaining the relative specificity for TAR over pre-miR-21 seen for Palbociclib (Supplementary Fig. 7). However, the B_max_ of this new compound for pre-miR-21 levels off at 0.4, suggesting the improved affinity is due to increased non-specific binding under low salt conditions. We conclude that it is not kinase binding that leads to high affinity and specific binding to RNA, but rather chemical features of the Palbociclib structure.

## Discussion

The discovery of drug-like small molecules that bind to RNA potently and specifically would open very broad new pharmaceutical opportunities^1–3^. However, reports of small molecules with potent (nM) and specific RNA binding activity are very few. Even successful programs leading to FDA approval^7^ have emerged from phenotypic screens with substantial uncertainties as to whether their mechanism of action is related to RNA, because binding to their putative RNA target is approximately 1,000-fold weaker than their cellular activity^8^. In the case of HIV TAR, by far the best studied model for small molecule binding to RNA, only a few molecules have been reported with the characteristics we observe for Palbociclib^17,18^, namely nM binding, specificity, and drug-like properties^40^, despite over 3 decades of investigations in academia and the pharmaceutical and biotech industries. As a result, it has been stated that targeting of RNA secondary structures with small molecules is unlikely to be successful, and HIV TAR itself was flagged as a typically challenging case^2^.

We show that Palbociclib and its derivative ‘Reverse Palbociclib’ specifically bind to HIV TAR with nM affinity under cellular ionic conditions and inhibit assembly of the Super Elongation Complex, which is essential for transactivation of transcription of the viral genome and the emergence of the virus from the latent state^16,38^. We hypothesize that this is possible because these molecules bind to both the apical loop and UCU bulge, as revealed by chemical shift mapping and initial investigation of the NOE data. In doing so, the small molecules create a new druggable binding pocket, as observed for the Hepatitis C IRES^26,27^ and our peptides^22,23^. This conclusion is supported by the unique NMR structural signature, distinct from both free TAR and from other complexes of TAR with peptides and small molecules; however, conclusive evidence for this latter hypothesis awaits formal structure determination, which is underway.

The new binding pocket provides for strong binding and specific inhibition of complex assembly in a rigorous biochemical assay with P-TEFb (Fig. 5). In contrast, cyclic peptides with 30 pM binding activity did not significantly affect SEC assembly on TAR and had disappointingly weak cellular activity^23^. We are very skeptical that numerous reports of antiviral activity associated with ligand binding to HIV TAR originated from on-target activity; indeed, the absence of such activity has been conclusively demonstrated for highly basic molecules such as aminoglycosides and basic peptoids, where antiviral activity was demonstrated to be unrelated to RNA binding^29,41^.

Palbociclib binds to HIV TAR with a KD of 100-300 nM, depending on ionic conditions, comparable to its reported affinity for cdk6 (K_D_ ~60nM)^24^, which was only achieved following a decade-long medicinal chemistry effort. Other studies have reported binding of FDA approved drugs to RNA^42–44^, including kinase inhibitors^45^, but binding in these cases was 100-1,000 fold weaker than we observe^25^ or non-specific^46,47^. It is sometimes stated that kinase inhibitors make good RNA binding molecules, but we have shown that RNA binding can be obtained fully independently of kinase inhibition. Indeed, Abemaciclib, a functionally related kinase inhibitor, showed only a weaker μM interaction. Additionally, by synthesizing ’Reverse Palbociclib’, we generated a molecule that binds to RNA better than Palbociclib, while not significantly inhibiting any kinase within a panel of more than 300 enzymes^39^.

Palbociclib inhibits HIV replication in cell lines that do not significantly express cdk4/cdk6^48^, but it would be very premature to attribute this inhibitory effect to direct binding to HIV TAR. The ‘Reverse Palbociclib’ structure was designed with the goal of addressing cellular activity by unambiguously separating kinase inhibition from RNA binding. Unfortunately, the ‘Reverse Palbociclib’ compound is toxic to multiple cell lines at single digit μM concentrations, for reasons we do not understand, preventing unbiased cellular investigations. It would be impossible to cleanly separate inhibition of viral replication from cellular toxicity for ‘Reverse Palbociclib’, and from kinase inhibition for Palbociclib itself. We are actively pursuing the synthesis of less toxic derivatives of this molecule that retain inhibitory activity.

The discovery that a blockbuster drug like Palbociclib binds to RNA with such potency and specificity demonstrates that drug-like molecules do indeed bind to and inhibit RNA, even in the absence of complex higher order RNA structures. Stem-loops distorted by internal loops and bulges, like HIV TAR, are the dominant secondary structure element in RNA and likely to form in cells where RNAs are less stably folded^14,12^, while more complex tertiary structures are unfolded by RNA-binding proteins, RNA helicases, and active translation. The present work demonstrates that drug-like small molecules not only bind to these functional RNAs but do so with the high affinity required for credible pre-clinical investigations.

## Methods

### RNA Transcription

All RNAs used for NMR were prepared *in house* using *in vitro* transcription on a large scale (typically 10 mL) using DNA oligonucleotide templates (IDT) and T7 RNA polymerase purified *in house*, essentially as described^49^. Briefly, 100 mL of 8 μM TOP DNA (5’-CTATAGTGAGTCGTATTA-3’), corresponding to the phage T7 RNA polymerase promoter region, was annealed to 80 μL of 100 μM template sequences with 13 mM MgCl_2_, heated to 95 °C for 4 min, then annealed at room temperature over 20 minutes. The annealing mixture was incubated with 5 mM of each of the four NTPs (ATP, GTP, UTP and CTP, from Sigma), 1x transcription buffer, 8% PEG-8000 and 35 mM magnesium chloride with 0.4 mg/mL T7 RNA polymerase.

All RNA samples are purified from crude transcriptions by 20% denaturing polyacrylamide gel electrophoresis (PAGE), electro-eluted and concentrated by ethanol precipitation. The samples were re-dissolved in ~12 mL of high salt wash (700 mM NaCl, 200 mM KCl, in 10 mM potassium phosphate at pH 6.5, with 10 μM EDTA to chelate any divalent ions) and concentrated using Centriprep conical concentrators (3,000 kDa cutoff, Millipore). The RNAs were then slowly exchanged into storage buffer (10 mM potassium phosphate at pH 6.5, with 10 mM NaCl and 10 μM EDTA). Prior to NMR experiments, all RNA samples were desalted using NAP-10 gravity columns, lyophilized, and re-dissolved in buffer, then annealed by heating for 4 minutes to 90 °C and snap cooling at −20 °C. For experiments investigating non-exchangeable protons, samples were lyophilized to dryness and dissolved into 99.99% D2O; samples used to study exchangeable protons were dissolved instead in 95% H_2_O/5% D_2_O.

### Fluorescence binding assays

Fluorescence binding assays were conducted in 50 mM bis-Tris buffer at pH 6.5, using a published method based on the incorporation of a single fluorescent base analogue (2-amino-purine, 2-AP) in place of U25, and by then monitoring changes in the reporter emission fluorescence intensity at 362 nm during small molecule titrations^37^. Each experiment was performed in duplicate, with 50 nM 2-AP-TAR (purchased from IDT) titrated with 5 mL of ligand stock solutions prepared by serial dilution. The previously reported method was modified to include a 250-fold excess yeast tRNA to the buffer, as a competitor, to reduce non-specific binding. Data were collected on a Horiba FL3-21tau Fluorescence Spectrophotometer, exported to Graphpad Prism v.8.1.1 and fit to equation 1,

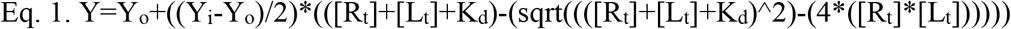

where *Y* is the measured fluorescence at 362 nm, *Y*_*i*_ is fluorescence of the fully bound RNA, *Y*_*o*_ is the fluorescence of the unbound RNA, *R*_*t*_ is total RNA concentration, *L*_*t*_ is total ligand concentration and *K*_*d*_ is the binding affinity.

### NMR binding assays

In order to identify small molecules that bind to RNA and quantify the strength of the interaction, we used NMR ligand-detected methods^33^. Compounds were first dissolved to 1-10 mM in pure water or DMSO, depending on their solubility, then prepared to 100 μM in 490 μL in buffer (50 mM d19-deuterated bis-Tris buffer at pH 6.5 containing 11.1 μM sodium 4,4-dimethyl-4-silapentane-1-sulfonate (DSA) as chemical shift and intensity reference, all dissolved in 99.99% D_2_O). The internal DSA reference allows us to directly compare the spectra (in the absence or presence of RNA) to identify binding by decreases in the NMR signals of the ligands due to increased rotational correlation time of the ligand when it binds to RNA^31^. The 9 protons on the internal reference also provide a control for ligand concentration. All 1D-^1^H NMR ligand-detected experiments were conducted using excitation sculpting water suppression (zgesgp, Bruker); a free ligand reference spectrum was collected, followed by titration in 10 μL steps of 500 μM RNA stock solution (10 μM final RNA concentration in the NMR tube). Each experiment was collected with 16 scans, 16k data points, and with a recycle delay of 1.0 sec, for an overall acquisition time of about 5 min per sample, including experimental setup.

We also generated titration curves in a higher salt buffer intended to mimic the conditions prevalent in the cell. A 10 mL stock of each compound at 100 μM concentration was dissolved in 50 mM d_19_-deuterated bis-Tris buffer, at pH 6.5, containing 11.1 μM DSA, plus 200 mM NaCl, 50 mM KCl and 4 mM MgCl_2_. Since these high salt conditions inevitably lead to RF heating of the sample, we increased the recycle delay from 1 to 5 sec. Compounds were divided into 12 1.5 mL microcentrifuge tubes at 490 μL each and titrated with 10 μL of RNA (0.5 to 1,000 μM). The final RNA concentration in each tube increased from 0.01 μM (10 nM) to 20 μM, while a tube with no RNA was used as control. Binding is detected by decreases in peak height relative to the free ligand over the concentration range of the added RNA. Affinity measurements were fit to single site binding curve models in Graphpad Prism v.8.1.1.

When titrating HIV TAR into a standard 100 μM sample of Palbociclib, the decrease in free ligand linewidth was plotted against RNA concentration. For proteins, which have a compact and globular shape, these curves can be directly fit using equation 2 below to extract apparent binding constants^31^. This approximation is also very likely to apply to an RNA the size of TAR, whose shape approximates an ellipsoid of rotation of aspect ratio close to 1.1-1.2, given its diameter of about 25 Å, accounting for hydration, and length of 30 Å. In the expression that follows, *I*_*B*_ is the intensity of the bound ligand peak height, *I*_*F*_ is the free ligand peak height, *P*_*t*_ is the total receptor concentration and *L*_*t*_ is the total ligand concentration. The constant c is the ratio of the bound peak width *ν*_*B*_ and the free peak width *ν*_*F*_, as shown in eq. 3:

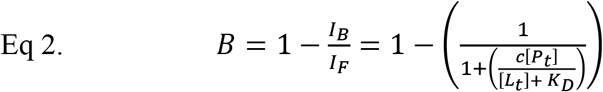

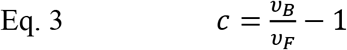

For proteins, the bound peak width *(ν*_*B*_) can be approximated by the molecular weight of the protein multiplied by a shape-related constant *ρ* as follows^31^:

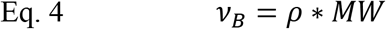

The quantitative analysis of equation 2-4 might not be fully warranted when a ligand induces large changes in target structure and shape upon binding; under these circumstances, variations in the line width constant *c* might not reliably allow measurement of bound ligand line width *ν*_*B*_, because it would change between free and bound ligand state. Changes in hydrodynamic shape are not likely to be large given the small size of TAR and its nearly spherical shape, but changes in this constant would nonetheless lead to uncertainties in the binding affinities. Therefore, plotting changes in ligand peak height vs RNA concentration only provides a semi-quantitative estimate of affinity. With this caveat, we fit the total change in ligand line width against RNA concentration using equation 5, where *B*_*max*_ is the maximum binding capacity and represents fully titrated or broadened ligand signal, *R*_*t*_ is the RNA concentration and *NS* is the slope of the nonlinear regression (non-specific binding is assumed to be linear with respect to RNA concentration):

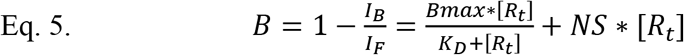

### RNA NMR analysis

RNAs are snap-cooled in 250-500 μL NMR buffer (50 mM d19 bis-Tris pH 6.5, 50 mM NaCl) by heating to 95 °C for 4 min then placed at −20 °C until frozen. Interactions were monitored through changes in chemical shifts in a series of NMR experiments, including 1D ^1^H, 2D ^1^H-^1^H TOCSY and ^1^H-^1^H NOESY experiments, which were all used to map the binding site of the ligand on the RNA. All pulse programs were standard sequences provided with the Bruker software package. Experiments in H_2_O NMR buffer were used to detect changes in exchangeable proton signals to assess the RNA secondary structure, while experiments in D_2_O NMR buffer were used to observe intermolecular NOEs between ligand and RNA.

After TAR RNA was fully titrated with a small molecule, initial RNA assignments for the complex were obtained by comparing TOCSY and NOESY spectra of the RNA:small molecule complex with those of free TAR, as reported by us in the past^50^, using well-established methods^49^.

### Molecular modeling

The core structure of ‘Reverse Palbociclib’ was derived through modeling in the Biosolveit SeeSAR package (v. 7) using the publicly available cdk6 enzyme bound to the three commercially available kinase inhibitors (Palbociclib, Abemaciclib and Ribociclib). The Biosolveit SeeSAR package was used to predict decreases in apparent affinity for cdk6, but not to interrogate affinity for the RNA, since the software has not been validated for work with RNA.

### Compound acquisition and synthesis

Commercially available small molecule compounds (Palbociclib, Abemaciclib and Ribociclib) were purchased from Selleckchem, while ‘Reverse Palbociclib’ was synthesized *in house* as follows (Supplementary Fig. 8).

#### Preparation of Intermediate-1

5-Amino-2-chloropyrimidine (1 equiv.), cyclopentanone (3 equiv.) and dichloromethane (0.2 M) were taken in a 100 mL round bottom flask (RBF) with a magnetic stir bar. The solution was cooled to 0 °C, then a solution of TiCl4 (1.2 equiv.) in 0.05 M of dichloromethane was added to the reaction mixture (RM) slowly over a period of 10 to 15 minutes. The RM was stirred at room temperature (RT) for 2 hours. Sodium cyanoborohydride (3 equiv.) was added in 4 equal portions over 10-minute time intervals and the RM stirred at RT for another 2 hours. The RM was cautiously quenched with water and extracted with ethyl acetate twice. The combined organic layer was dried over Na_2_SO_4_, filtered, and concentrated under vacuum. The crude RM was purified on silica gel using 0-30% ethyl acetate in hexanes as eluent. Relevant fractions were evaporated in vacuum to give *Intermediate-1* (Yield: 65%).

#### Preparation of Intermediate-2. Intermediate-1

(1 equiv.), iron powder (0.1 equiv.) and dichloromethane (0.2 M) were taken in a 100 mL RBF with a magnetic stir bar. The solution was cooled to 0 °C, then a solution of Br_2_ (1.2 equiv.) in 0.05 M of dichloromethane was added to the reaction mixture slowly over a period of 10 to 15 minutes. The RM was stirred at RT overnight (20-24 hours). The iron powder was filtered off, washed with small amount of dichloromethane and the filtrate was concentrated under vacuum. The crude RM was purified on silica gel using 0-20% ethyl acetate in hexanes as eluent. Relevant pure fractions were evaporated in vacuum to give *Intermediate-2* (Yield: 70%).

#### Preparation of Intermediate-3. Intermediate-2

(1 equiv.), (Z)-(4-Ethoxy-4-oxo-2-buten-2-yl) boronic acid pinacol ester (1.2 equiv.), Na_2_CO_3_ (2.7 equiv.) and a mixture of DMF/H_2_O (5:1, 0.25 M) were taken in a microwave vial with a magnetic stir bar. The solution was purged with N2 gas for 10-15 minutes, then catalyst PdCl_2_ (PPh_3_)_2_ (0.05 equiv.) was added to the RM and purged the N_2_ gas for another 2 to 5 minutes. The RM was sealed and stirred at 90 °C overnight (18-23 hours). The RM was cooled to room temperature, filtered off and the filtrate was diluted with ethyl acetate, and washed with brine solution. The organic layer was dried over Na_2_SO_4_, filtered, and concentrated under vacuum. The crude RM was purified on silica gel using 0-30% ethyl acetate in hexanes as eluent. Relevant pure fractions were evaporated under vacuum to give *Intermediate-3* (Yield: 48%).

#### Preparation of Intermediate-4

Int-3 (1 equiv.), Cs_2_CO_3_ (1.2 equiv.) and DMF (0.25 M) were taken in a 100 ml RBF with a magnetic stir bar. The reaction mixture was stirred at room temperature overnight (20-24 hours). The RM was diluted with ethyl acetate, then washed with brine solution twice. The organic layer was dried over Na_2_SO_4_ and concentrated under vacuum. The crude RM was purified on silica gel using 0-30% ethyl acetate in hexanes as eluent. Relevant fractions were evaporated under vacuum to give *Intermediate-4* (Yield: 50%).

#### Preparation of ‘Reverse Palbociclib’

A 2-necked RBF was purged and maintained under an atmosphere of nitrogen. The corresponding aniline (2.1 equiv.) and toluene (0.20 M) were taken in an RBF with a magnetic stir bar. The RM was cooled to 0 °C, then LIHMDS (1 M solution in THF, 2.1 equiv.) was added to the RM over period of 2-5 minutes. After 5-10 minutes, a solution of the corresponding *Intermediate-4* (1 equiv.) in 0.05 M of toluene was added to the RM at 0 °C. The RM was then stirred at room temperature for 2-4 hours. The RM was quenched with aqueous saturated NaHCO_3_ solution and diluted with ethyl acetate. The organic layer was dried over Na_2_SO_4_ and concentrated under vacuum. The crude RM was purified on silica gel using 0-80% ethyl acetate in hexanes as eluent. Relevant pure fractions were evaporated under vacuum to give *Intermediate-5*, which was dissolved in DCM/TFA (4:1, 0.05 M), then stirred for 1-2 hours at RT. The crude reaction was concentrated and purified on HPLC using water/acetonitrile as eluent. Relevant pure peak fractions were lyophilized to give the corresponding final compound *Reverse Palbociclib* (overall yield for two steps: 35%).

### Kinase inhibition

Kinase inhibition profiling was conducted at *Nanosyn* using the Nanosyn Caliper profiling system at 1 and 10 μM concentration of small molecule, to profile a total of 330 kinases, following published methods^39^.

### PTEFb-binding assay

The P-TEFb/Tat1:57/AFF4 binding assays were run using EMSA as described^23,38^, with protein samples generously donated by Drs. Ursula Schulze-Gahmen and James Hurley, except that binding buffers also contained 250-fold excess of yeast tRNA to reduce non-specific binding. Palbociclib-bound samples were added in 20-fold excess over ^32^P labeled TAR (10 nM Palbociclib, 0.5 nM ^32^P TAR) and the ligand-RNA complex was equilibrated for 30 min at room temperature prior to titration with the pre-formed P-TEFb/Tat1:57/AFF4 complex.

## Acknowledgements

We wish to thank all members of the Varani group for discussion and support; Dr. Greg Olsen for help with final preparation of the manuscript; Drs. Ursula Schulze-Gahmen and James Hurley for donation of P-TEFb samples; this project was funded by grant NIH NIGMS 1R35GM126942 to GV.

## Author Contributions

M.D.S. collected and analyzed the data; V.V. performed compound synthesis. The research was conceived by

M.D.S. and G.V. and supervised by G.V. The manuscript was written by M.D.S., V.V. and G.V. All authors have approved the final version of the manuscript.

## Additional information

Supplementary material is available: supplementary figures 1 – 8 and supplementary table 1.

## References

1. Hermann, T. Drugs targeting the ribosome. Current Opinion in Structural Biology 15, 355–366 (2005).

2. Warner, K. D., Hajdin, C. E. & Weeks, K. M. Principles for targeting RNA with drug-like small molecules. Nature Reviews Drug Discovery 17, 547–558, doi:10.1038/nrd.2018.93 (2018).

3. Gallego, J. & Varani, G. Targeting RNA with Small Molecule Drugs: Therapeutic Promises and Chemical Challenges. Acc. Chem. Res. 34, 836–843 (2001).

4. Rizvi, N. F. et al. Targeting RNA with Small Molecules: Identification of Selective, RNA-Binding Small Molecules Occupying Drug-Like Chemical Space. Slas Discovery 25, 384–396, doi: 10.1177/2472555219885373 (2020).

5. Rizvi, N. F. et al. Discovery of Selective RNA-Binding Small Molecules by Affinity Selection Mass Spectrometry. Acs Chemical Biology 13, 820–831, doi:10.1021/acschembio.7b01013 (2018).

6. Palacino, J. et al. SMN2 splice modulators enhance U1-pre-mRNA association and rescue SMA mice. Nature Chemical Biology 11, 511–517 (2015).

7. Ratni, H. et al. Discovery of Risdiplam, a Selective Survival of Motor Neuron-2 (SMN2) Gene Splicing Modifier for the Treatment of Spinal Muscular Atrophy (SMA). Journal of Medicinal Chemistry 61, 6501–6517, doi:10.1021/acs.jmedchem.8b00741 (2018).

8. Campagne, S. et al. Structural basis of a small molecule targeting RNA for a specific splicing correction. Nature Chem. Biol. 15, 1191–1198 (2019).

9. Shortridge, M. D. & Varani, G. Structure based approaches for targeting non-coding RNAs with small molecules. Current Opinion in Structural Biology 30, 79–88, doi:10.1016/j.sbi.2015.01.008 (2015).

10. Howe, J. A. et al. Selective small-molecule inhibition of an RNA structural element. Nature 526, 672–677 (2015).

11. Flynn, R. A. et al. Transcriptome-wide interrogation of RNA secondary structure in living cells with icSHAPE. Nature Protocols 11, 273–290 (2016).

12. Marinus, T., Fessler, A. B., Ogle, C. A. & Incarnato, D. A novel SHAPE reagent enables the analysis of RNA structure in living cells with unprecedented accuracy. Nucleic Acids Research 49, doi:10.1101/2020.08.31.274761 (2021).

13. Wu, X. B. & Bartel, D. P. Widespread Influence of 3’-End Structures on Mammalian mRNA Processing and Stability. Cell 169, 905–917, doi:10.1016/j.cell.2017.04.036 (2017).

14. Wan, Y. et al. Landscape and variation of RNA secondary structure across the human transcriptome. Nature 505, 706–709, doi:10.1038/nature12946 (2014).

15. Hewitt, W. M., Calabrese, D. R. & Schneekloth, J. S. Evidence for ligandable sites in structured RNA throughout the Protein Data Bank. Bioorganic & Medicinal Chemistry 27, 2253–2260, doi:10.1016/j.bmc.2019.04.010 (2019).

16. Karn, J. Tackling Tat. J. Mol. Biol. 293, 235–254 (1999).

17. Murchie, A. I. H. et al. Structure-based Drug Design Targeting an Inactive RNA Conformation: Exploiting the Flexibility of HIV-1 TAR RNA. J. Mol. Biol. 336, 625–638 (2004).

18. Davis, B. et al. Rational Design of Inhibitors of HIV-1 TAR RNA through the Stabilisation of Electrostatic “Hot Spots”. J. Mol. Biol. 336, 343–356 (2004).

19. Mei, H.-Y. et al. Discovery of Selective, Small-molecule Inhibitors of RNA Complexes -I. The Tat Protein/TAR RNA Complexes Required for HIV-1 Transcription. Bioorg. Med. Chem. 5, 1173–1184 (1997).

20. Mei, H.-Y. et al. Inhibitors of Protein-RNA Complexation that Target the RNA: Specific Recognition of Human Immunodeficiency Virus Type I TAR RNA by Small Organic Molecules. Biochemistry 37, 14204–14212 (1998).

21. Lalonde, M. S. et al. Inhibition of Both HIV-1 Reverse Transcription and Gene Expression by a Cyclic Peptide that Binds the Tat-Transactivating Response Element (TAR) RNA. Plos Pathogens 7, e1002038 (2011).

22. Davidson, A. et al. Simultaneous recognition of HIV-1 TAR RNA bulge and loop sequences by cyclic peptide mimics of Tat protein Proc. Natl. Acad. Sci. USA 106, 11931–11936 (2009).

23. Shortridge, M. D. et al. An ultra-high affinity ligand of HIV-1 TAR reveals the RNA structure recognized by P-TEFb. Nucleic Acids Res 47, 1523–1531, doi:10.1093/nar/gky1197 (2018).

24. Toogood, P. L. et al. Discovery of a Potent and Selective Inhibitor of Cyclin-Dependent Kinase 4/6. Journal of Medicinal Chemistry 48, 2388–2406, doi:10.1021/jm049354h (2005).

25. Haniff, H. S. et al. Design of a small molecule that stimulates vascular endothelial growth factor A enabled by screening RNA fold-small molecule interactions. Nature Chemistry 12, 952–961, doi:10.1038/s41557-020-0514-4 (2020).

26. Parsons, J. et al. Conformational inhibition of the hepatitis C virus internal ribosome entry site RNA. Nature Chemical Biology 5, 823–825 (2009).

27. Dibrov, S. M. et al. Structure of a hepatitis C virus RNA domain in complex with a translation inhibitor reveals a binding mode reminiscent of riboswitches. Proceedings of the National Academy of Sciences of the United States of America 109, 5223–5228, doi:10.1073/pnas.1118699109 (2012).

28. Hamy, F. et al. A New Class of HIV-1 Tat Antagonist Acting through Tat-TAR Inhibition. Biochemistry 37, 5086–5095 (1998).

29. Hamy, F. et al. An Inhibitor of the Tat/TAR RNA Interaction that Effectively Suppresses HIV-1 Replication. Proc. Natl. Acad. Sci. USA 94, 3548–3553 (1997).

30. Zeiger, M. et al. Fragment based search for small molecule inhibitors of HIV-1 Tat-TAR. Bioorganic & Medicinal Chemistry Letters 24, 5576–5580 (2014).

31. Shortridge, M. D., Hage, D. S., Harbison, G. S. & Powers, R. Estimating Protein’s Ligand Binding Affinity Using High-Throughput Screening by NMR. Journal of Combinatorial Chemistry 10, 948–958 (2008).

32. Abulwerdi, F. A. et al. Development of Small Molecules with a Noncanonical Binding Mode to HIV-1 Trans Activation Response (TAR) RNA. Journal of Medicinal Chemistry 59, 11148–11160 (2016).

33. Shortridge, M. D. & Varani, G. Efficient NMR Screening Approach to Discover Small Molecule Fragments Binding Structured RNA. Acs Medicinal Chemistry Letters 12, 1253–1260, doi:10.1021/acsmedchemlett.1c00109 (2021).

34. Dalvit, C., Fogliatto, G. P., Stewart, A., Veronesi, M. & Stockman, B. WaterLOGSY as a metod for primary NMR screening: Practical aspects and range of applicability. J. Biomol. NMR 21, 349–359 (2001).

35. Mayer, M. & Mayer, B. Group Epitope Mapping by Saturation Transfer Difference NMR To Identify Segments of a Ligand in Direct Contact with a Protein Receptor. J. Am. Chem. Soc. 123, 6108–6117 (2001).

36. Mayer, M. & James, T. Detecting ligand binding to a small RNA target via saturation transfer difference NMR experiments in D2O and H2O. Journal of the American Chemical Society 124, 13376–13377 (2002).

37. Bradrick, T. D. & Marino, J. P. Ligand-Induced Changes in 2-Aminopurine Flourescence as a probe for Small Molecule Binding to HIV-1 TAR RNA. RNA 10, 1459–1468 (2004).

38. Schulze-Gahmen, U. et al. Insights into HIV-1 proviral transcription from integrative structure and dynamics of the Tat:AFF4:P-TEFb:TAR complex. eLife 5, e15910, doi:10.7554/eLife.15910 (2016).

39. Elkins, J. M. et al. Comprehensive characterization of the Published Kinase Inhibitor Set. Nature Biotechnology 34, 95–103, doi:10.1038/nbt.3374 (2016).

40. Lipinski, C. A., Lombardo, F., Dominy, B. W. & Feeney, P. J. Experimental and Computational Approaches to Estimate Solubility and Permeability in Drug Discovery and Development Setting. Adv. Drug Deliv. Rev. 23, 3–25 (1997).

41. Daelemans, D. et al. A second target for the peptoid Tat/transactivation response element inhibitor CGP64222: inhibition of human immunodeficiency virus replication by blocking CXC-chemokine receptor 4-mediated virus entry. Mol Pharmacol 57, 116–124 (2000).

42. Zheng, S., Chen, Y., Donahue, C. P., Wolfe, M., s. & Varani, G. Structural Basis for Stabilization of the Tau Pre-mRNA Splicing Regulatory Element by Novantrone (Mitoxantrone). Chemistry & Biology 16, 557–566, doi:doi:10.1016/j.chembiol.2009.03.009 (2009).

43. LeBlanc, R. M. et al. Structural insights of the conserved "priming loop" of hepatitis B virus pre-genomic RNA. J Biomol Struct Dyn, 1–13, doi:10.1080/07391102.2021.1934544 (2021).

44. Chen, J. L. et al. Design, Optimization, and Study of Small Molecules That Target Tau Pre-mRNA and Affect Splicing. Journal of the American Chemical Society 142, 8706–8727, doi:10.1021/jacs.0c00768 (2020).

45. Zhang, P. Y. et al. Reprogramming of Protein-Targeted Small-Molecule Medicines to RNA by Ribonuclease Recruitment. Journal of the American Chemical Society 143, 13044–13055, doi:10.1021/jacs.1c02248 (2021).

46. Liu, Y. et al. Mitoxantrone analogues as ligands for a stem-loop structure of tau pre-mRNA. Journal of medicinal chemistry 52, 6523–6526, doi:10.1021/jm9013407 (2009).

47. Velagapudi, S. P. et al. Approved Anti-cancer Drugs Target Oncogenic Non-coding RNAs. Cell Chemical Biology 25, 1086–1094, doi:DOI 10.1016/j.chembiol.2018.05.015 (2018).

48. Pauls, E. et al. Palbociclib, a selective inhibitor of cyclin-dependent kinase4/6, blocks HIV-1 reverse transcription through the control of sterile alpha motif and HD domain-containing protein-1 (SAMHD1) activity. Aids 28, 2213–2222 (2014).

49. Varani, G., Aboul-ela, F. & Allain, F. H.-T. NMR Investigations of RNA Structure. Progr. NMR Spectr. 29, 51–127 (1996).

50. Aboul-ela, F., Karn, J. & Varani, G. Structure of HIV-1 TAR RNA in the Absence of Ligands Reveals a Novel Conformation of the Trinucleotide Bulge. Nucleic Acids Res. 24, 3974–3981 (1996).

