## Supplementary material for "The kinase inhibitor Palbociclib is a potent and specific RNA-binding molecule"

**Supplementary figures 1 – 8, and supplementary table 1.**

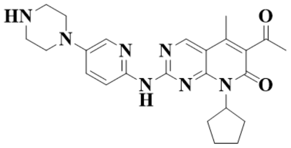

**Palbociclib**

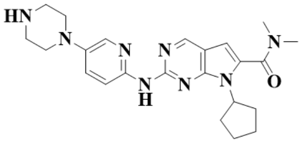

**Ribociclib**

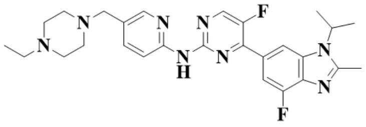

**Abemaciclib**

**Supplementary Fig. 1. Chemical structures of the three FDA-approved cdk4/cdk6 drugs, Palbociclib, Ribociclib and Abemaciclib.**

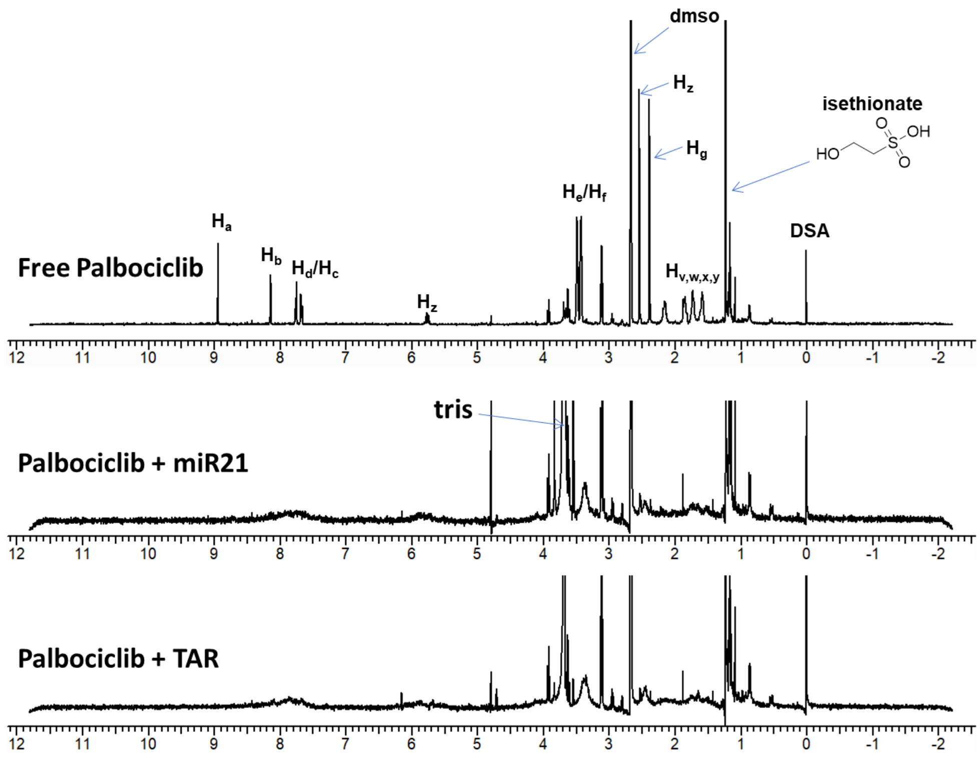


**Supplementary Fig. 2. At 10 µM concentrations, Palbociclib binds to both HIV TAR RNA and to pre-miR-21.** Top, reference spectrum of 100 µM free ligand; Middle, spectrum of Palbociclib following addition of 10 µM pre-miR-21 to the free ligand; Bottom, spectrum of Palbociclib following addition of 10 µM HIV TAR to the free ligand. These spectra were recorded in deuterated 50 mM Bis-Tris buffer (pH 6.5) without any added counterions. The ligand signals and other buffer components are labeled in the spectra. The decrease in ligand signal is due to the increase in rotational correlation time ($\tau_c$) of the ligand when bound to the RNA. Compounds that do not bind (e.g. DSA) do not show a decrease in the NMR signal. The broad weak humps around 8 and 5.5 ppm represent the RNA signal; the top spectrum is shown at a reduced noise level to emphasize the disappearance of the Palbociclib NMR signals.

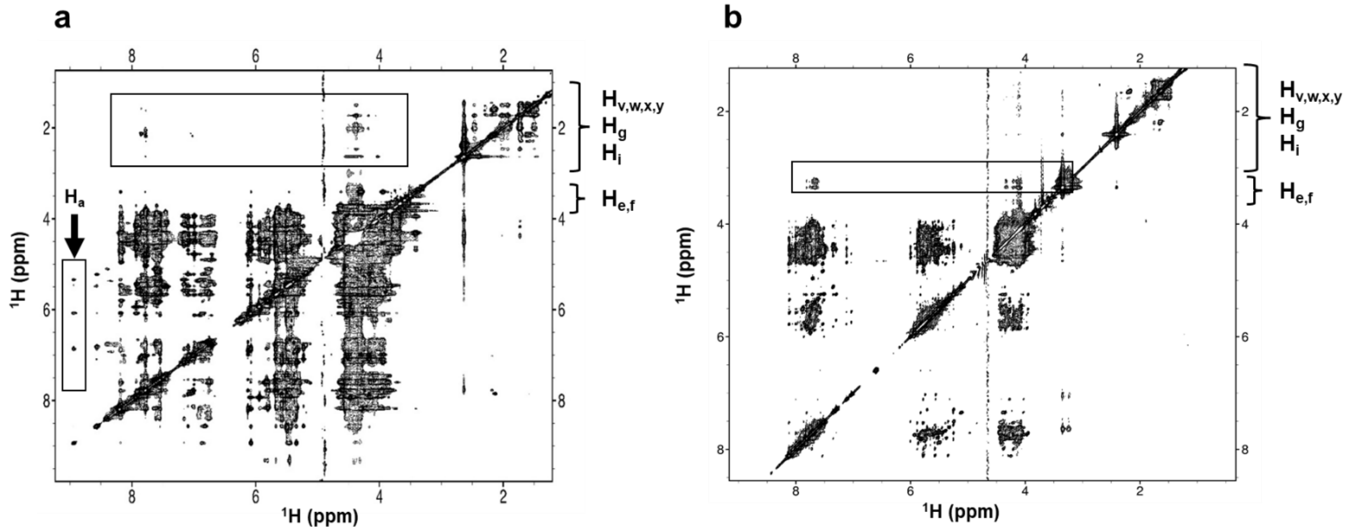


**Supplementary Fig. 3. NOESY spectra of Palbociclib with TAR and pre-miR-21 have very different characteristics.** a) Many intermolecular NOE interactions are observed between HIV TAR and Palbociclib. b) In contrast, pre-miR-21 shows a different structural fingerprint, with markedly fewer NOEs, consistent with µM binding and a dynamic, poorly defined binding site. Each spectrum was collected at 800 MHz, with a 300 ms mixing time; the samples contained 500 µM RNA and 750 µM Palbociclib, in 50 mM d9-bisTris pH 6.5, 50 mM NaCl, in 99.99% D2O.

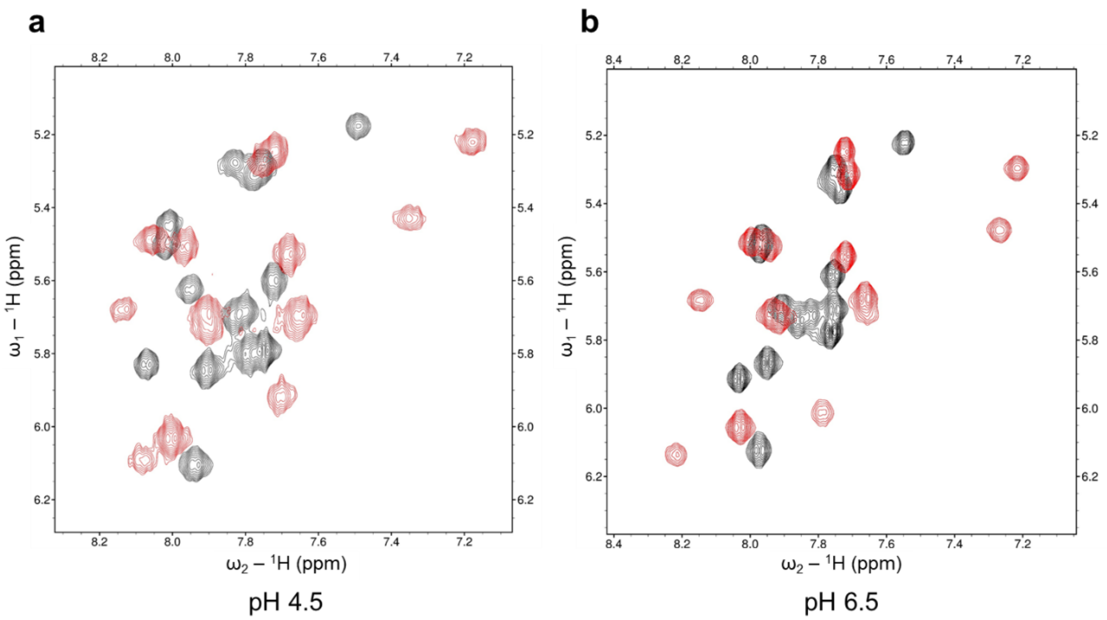


**Supplementary Fig. 4. Binding of Palbociclib to HIV TAR under different pH conditions.** a) Sample prepared in 50 mM Sodium Acetate at pH 4.5, and b) sample prepared in 50 mM bis-Tris at pH 6.5. All TOCSY spectra were collected at 800 MHz under high salt conditions (200 mM NaCl, 50 mM KCl and 4 mM MgCl2) at 37 °C. 250 µM RNA (black) was titrated with Palbociclib (red) until saturation was reached (as established from the absence of further changes in the spectra) and changes in chemical shifts were recorded.

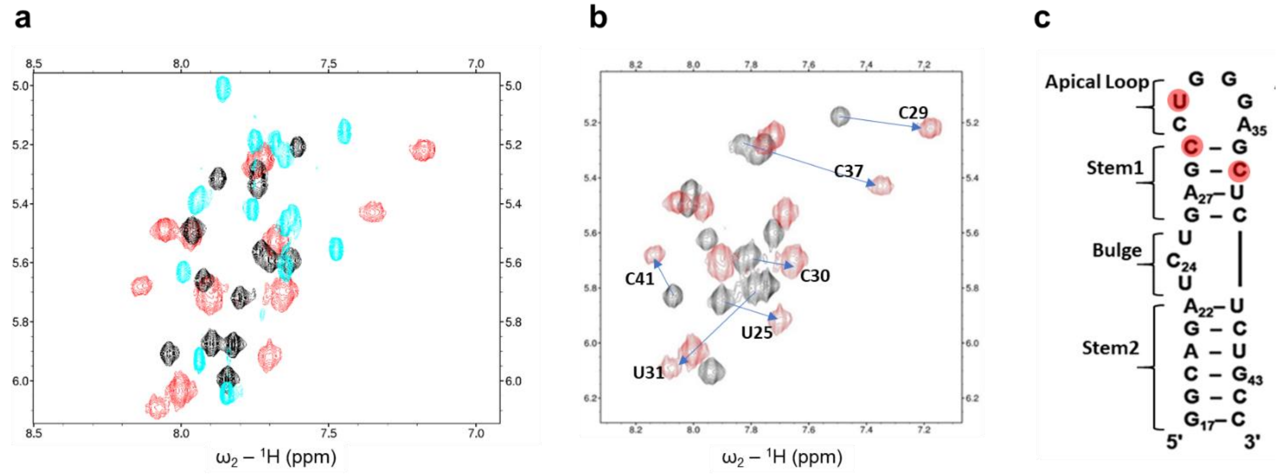


**Supplementary Fig. 5. Unique structural response to Palbociclib binding.** TOCSY spectra provide a structural fingerprint of ligand binding**.** (a) The large chemical shift changes between free (black) and bound (red) spectra, coupled with the large chemical shift changes between the JB181 (cyan) and Palbociclib bound spectra (red) suggest Palbociclib induces a third, so far uncharacterized structure of HIV TAR, distinct both from ligand-free TAR[50] and the structure bound to JB-181[23]. (b, c) While spectral changes are widespread, the largest chemical shift changes occur in the apical loop (nucleotides C29, U31 and C37), suggesting Palbociclib binds in the RNA major groove and can bridge the UCU bulge and RNA apical loop, as we observed for peptide JB181[23].

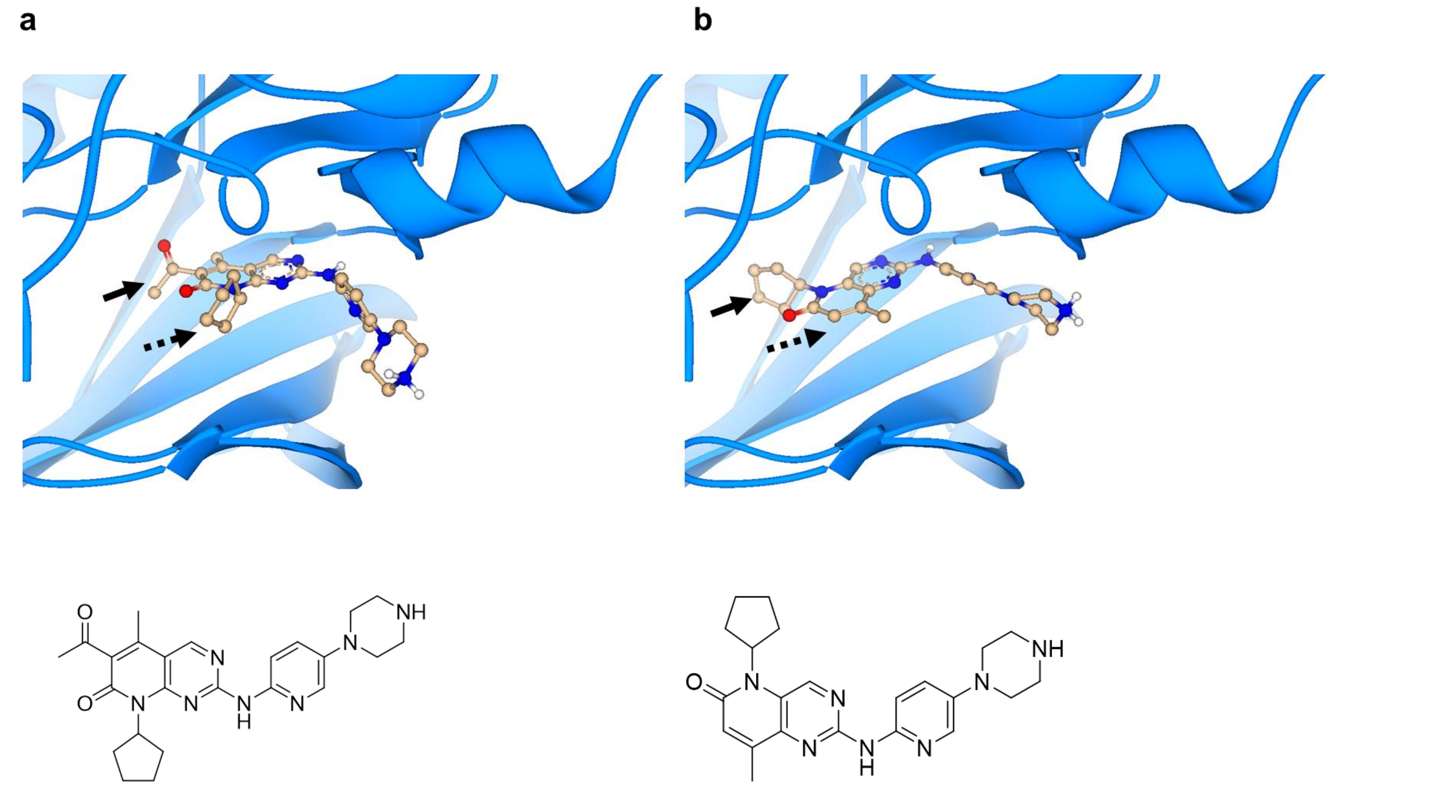


**Supplementary Fig. 6. Comparison of the cdk6 complexes of Palbociclib and a model for 'Reverse Palbociclib', bound to the catalytic site of the kinase.** a) The cyclopentane ring of Palbociclib binds in the hydrophobic pocket created by the flexible loop region of the kinase (dashed arrow) (PDB 5L2I). b) Predicted pose for 'Reverse Palbociclib' in the Palbociclib binding site. In these docking studies, the bridge NH between the pyrimidine and the pyridine attempts to maintain the hydrogen bond with the backbone of Val101, but the Pyrido[3,2,D]pyrimidin-6-one core structure of 'Reverse Palbociclib' flips the cyclopentane ring to occupy the pocket typically filled with the Palbociclib acetate group (solid arrow). This 'flip' reduces predicted affinity for cdk6 by >4 orders of magnitude. This hypothesis is supported by the data of Supplementary Table 1 demonstrating lack of inhibition of a panel of 330 kinases. The acetate group of Palbociclib was removed in 'Reverse Palbociclib', to improve synthetic accessibility.

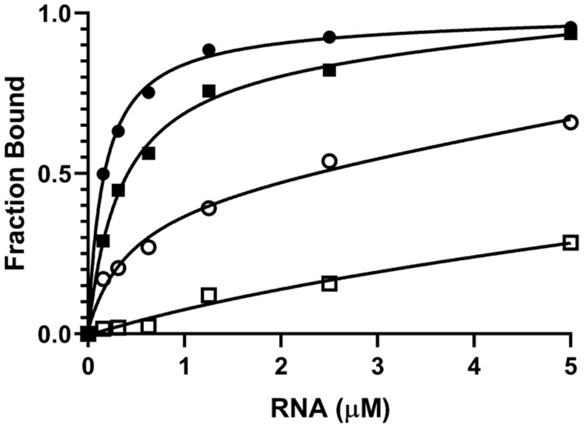


**Supplementary Fig. 7. Reverse Palbociclib shows improved binding affinity for HIV TAR and retains specificity over pre-miR-21.** NMR binding affinities for Palbociclib (squares) bound to HIV TAR (■) were fit to an apparent KD=300 nM and, for pre-miR-21 (□) to a KD=4,400 nM, under 'high salt' conditions (200mM NaCl, 50mM KCl and 4mM MgCl2). The Pyrido[3,2,D]pyrimidin-6-one core structure of Reverse Palbociclib provides improved binding affinity for HIV TAR, with resulting affinities improved to 100 nM for HIV TAR (●) and to 400 nM for pre-miR-21 (○). The Bmax values also increased for both RNAs, indicating reduced non-specific binding to RNA (compare squares and circles); this likely originates from the increased ionic strength conditions used for this titration, which would undoubtedly reduce non-specific binding due to the charge associated with the weakly basic piperazine group common to both molecules (predicted pKa 8-8.5).

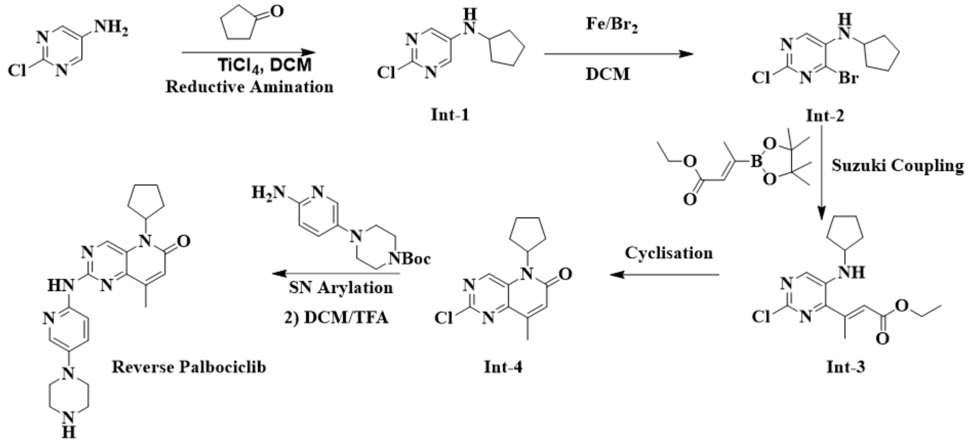


**Supplementary Fig. 8**. Synthetic scheme for the synthesis of 'Reverse Palbociclib'.

**Supplementary Table S1**: Summary of the kinome-wide investigation of kinase inhibition by 'Reverse Palbociclib', executed using the Nanosym Caliper technology[39]. The first column is the kinase identifier; the second column compound concentration and the third column % inhibition at the concentration tested.

| Kinase | Conc. Tested (μM) | Compound 54 |
|---|---|---|
| ABL1 | 10 | 3 |
| ABL1 | 1 | -2 |
| AKT1 | 10 | -2 |
| AKT1 | 1 | -4 |
| AKT2 | 10 | 3 |
| AKT2 | 1 | -6 |
| AKT3 | 10 | 1 |
| AKT3 | 1 | 1 |
| ALK | 10 | 0 |
| ALK | 1 | -3 |
| ALK2 | 10 | -4 |
| ALK2 | 1 | 4 |
| ALK4 | 10 | 6 |
| ALK4 | 1 | 6 |
| ALK5 | 10 | 15 |
| ALK5 | 1 | 9 |
| ALK6 | 10 | -12 |
| ALK6 | 1 | -4 |
| AMPK-A1B1G1 | 10 | 5 |
| AMPK-A1B1G1 | 1 | -2 |
| AMPK-A1B2G1 | 10 | 6 |
| AMPK-A1B2G1 | 1 | -10 |
| AMPK-A2B1G1 | 10 | 7 |
| AMPK-A2B1G1 | 1 | -3 |
| AMPK-A2B2G1 | 10 | 6 |
| AMPK-A2B2G1 | 1 | 2 |
| ARG | 10 | 7 |
| ARG | 1 | -2 |
| ARK5 | 10 | 1 |
| ARK5 | 1 | -6 |
| ASK1 | 10 | -2 |
| ASK1 | 1 | 0 |
| ATM | 10 | 41 |
| ATM | 1 | 21 |
| ATR | 10 | 55 |
| ATR | 1 | 29 |
| AURORA-A | 10 | 6 |
| AURORA-A | 1 | 0 |
| AURORA-B | 10 | 49 |

| | | |
|---|---|---|
| **AURORA-B** | **1** | 4 |
| **AURORA-C** | **10** | 7 |
| **AURORA-C** | **1** | 5 |
| **AXL** | **10** | 0 |
| **AXL** | **1** | -5 |
| **BLK** | **10** | -1 |
| **BLK** | **1** | -8 |
| **BMPR2** | **10** | -3 |
| **BMPR2** | **1** | 0 |
| **BMX** | **10** | 2 |
| **BMX** | **1** | -3 |
| **BRAF** | **10** | 8 |
| **BRAF** | **1** | 4 |
| **BRK** | **10** | -5 |
| **BRK** | **1** | -10 |
| **BRSK1** | **10** | 8 |
| **BRSK1** | **1** | -7 |
| **BRSK2** | **10** | 1 |
| **BRSK2** | **1** | -5 |
| **BTK** | **10** | -9 |
| **BTK** | **1** | -11 |
| **BUB1** | **10** | 66 |
| **BUB1** | **1** | 19 |
| **CAMK1A** | **10** | 4 |
| **CAMK1A** | **1** | 0 |
| **CAMK1D** | **10** | 3 |
| **CAMK1D** | **1** | -2 |
| **CAMK2A** | **10** | 1 |
| **CAMK2A** | **1** | -1 |
| **CAMK2B** | **10** | 3 |
| **CAMK2B** | **1** | 0 |
| **CAMK2D** | **10** | 2 |
| **CAMK2D** | **1** | -2 |
| **CAMK2G** | **10** | 8 |
| **CAMK2G** | **1** | -4 |
| **CAMK4** | **10** | -6 |
| **CAMK4** | **1** | -2 |
| **CAMKK1** | **10** | 0 |
| **CAMKK1** | **1** | -4 |
| **CDK1-CYCLINB** | **10** | -8 |
| **CDK1-CYCLINB** | **1** | -13 |
| **CDK2-CYCLINA** | **10** | 13 |
| **CDK2-CYCLINA** | **1** | 2 |
| **CDK2-CYCLINE** | **10** | 3 |
| **CDK2-CYCLINE** | **1** | 0 |
| **CDK3-CYCLINE** | **10** | 4 |

| | | |
|---|---|---|
| **CDK3-CYCLINE** | **1** | 1 |
| **CDK4-CYCLIND** | **10** | -6 |
| **CDK4-CYCLIND** | **1** | -4 |
| **CDK5-P25** | **10** | -9 |
| **CDK5-P25** | **1** | -8 |
| **CDK5-P35** | **10** | 5 |
| **CDK5-P35** | **1** | 2 |
| **CDK6-CYCLIND3** | **10** | -6 |
| **CDK6-CYCLIND3** | **1** | -12 |
| **CDK7** | **10** | -6 |
| **CDK7** | **1** | -2 |
| **CDK9-CYCLINT1** | **10** | -5 |
| **CDK9-CYCLINT1** | **1** | -3 |
| **CHEK1** | **10** | 3 |
| **CHEK1** | **1** | 0 |
| **CHEK2** | **10** | 4 |
| **CHEK2** | **1** | 0 |
| **CK1** | **10** | 63 |
| **CK1** | **1** | 9 |
| **CK1-DELTA** | **10** | 81 |
| **CK1-DELTA** | **1** | 23 |
| **CK1-EPSILON** | **10** | 41 |
| **CK1-EPSILON** | **1** | 1 |
| **CK1-GAMMA1** | **10** | -33 |
| **CK1-GAMMA1** | **1** | -14 |
| **CK1-GAMMA2** | **10** | -12 |
| **CK1-GAMMA2** | **1** | -12 |
| **CK1-GAMMA3** | **10** | -10 |
| **CK1-GAMMA3** | **1** | -8 |
| **CK2** | **10** | -12 |
| **CK2** | **1** | -13 |
| **CK2A2** | **10** | 1 |
| **CK2A2** | **1** | 4 |
| **CLK1** | **10** | 35 |
| **CLK1** | **1** | 2 |
| **CLK2** | **10** | 2 |
| **CLK2** | **1** | -13 |
| **CLK3** | **10** | -28 |
| **CLK3** | **1** | -4 |
| **CLK4** | **10** | 78 |
| **CLK4** | **1** | 18 |
| **CRAF** | **10** | -19 |
| **CRAF** | **1** | -8 |
| **CRIK** | **10** | -1 |
| **CRIK** | **1** | 7 |
| **CSK** | **10** | 2 |

| | | |
|---|---|---|
| **CSK** | **1** | -8 |
| **DAPK1** | **10** | 3 |
| **DAPK1** | **1** | 3 |
| **DAPK3** | **10** | -3 |
| **DAPK3** | **1** | -8 |
| **DCAMKL1** | **10** | 5 |
| **DCAMKL1** | **1** | -13 |
| **DCAMKL2** | **10** | -4 |
| **DCAMKL2** | **1** | -8 |
| **DDR1** | **10** | 7 |
| **DDR1** | **1** | 3 |
| **DDR2** | **10** | 0 |
| **DDR2** | **1** | -11 |
| **DGKa** | **10** | 7 |
| **DGKa** | **1** | -10 |
| **DGKb** | **10** | -2 |
| **DGKb** | **1** | -4 |
| **DGKz** | **10** | -49 |
| **DGKz** | **1** | -30 |
| **DLK** | **10** | 3 |
| **DLK** | **1** | 1 |
| **DRAK1** | **10** | 0 |
| **DRAK1** | **1** | -7 |
| **DRAK2** | **10** | -10 |
| **DRAK2** | **1** | -4 |
| **DYRK1A** | **10** | 1 |
| **DYRK1A** | **1** | -10 |
| **DYRK1B** | **10** | 25 |
| **DYRK1B** | **1** | 3 |
| **DYRK2** | **10** | 15 |
| **DYRK2** | **1** | -3 |
| **DYRK3** | **10** | 7 |
| **DYRK3** | **1** | -7 |
| **DYRK4** | **10** | -2 |
| **DYRK4** | **1** | -3 |
| **EGFR** | **10** | 24 |
| **EGFR** | **1** | -6 |
| **EIF2AK1** | **10** | 8 |
| **EIF2AK1** | **1** | -1 |
| **EIF2AK2** | **10** | -1 |
| **EIF2AK2** | **1** | -14 |
| **EIF2AK3** | **10** | 10 |
| **EIF2AK3** | **1** | -7 |
| **EIF2AK4** | **10** | 15 |
| **EIF2AK4** | **1** | 4 |
| **EPH-A1** | **10** | 0 |

| | | |
|---|---|---|
| **EPH-A1** | **1** | -4 |
| **EPH-A2** | **10** | 15 |
| **EPH-A2** | **1** | 9 |
| **EPH-A3** | **10** | 6 |
| **EPH-A3** | **1** | 8 |
| **EPH-A4** | **10** | 1 |
| **EPH-A4** | **1** | -2 |
| **EPH-A5** | **10** | 7 |
| **EPH-A5** | **1** | 2 |
| **EPH-A6** | **10** | 3 |
| **EPH-A6** | **1** | -1 |
| **EPH-A7** | **10** | -2 |
| **EPH-A7** | **1** | -2 |
| **EPH-A8** | **10** | 2 |
| **EPH-A8** | **1** | -3 |
| **EPH-B1** | **10** | 1 |
| **EPH-B1** | **1** | -4 |
| **EPH-B2** | **10** | -11 |
| **EPH-B2** | **1** | -30 |
| **EPH-B3** | **10** | 1 |
| **EPH-B3** | **1** | -2 |
| **EPH-B4** | **10** | -1 |
| **EPH-B4** | **1** | -6 |
| **ERB-B2** | **10** | 24 |
| **ERB-B2** | **1** | -6 |
| **ERB-B4** | **10** | -5 |
| **ERB-B4** | **1** | -7 |
| **FAK** | **10** | 9 |
| **FAK** | **1** | 6 |
| **FER** | **10** | 37 |
| **FER** | **1** | 7 |
| **FES** | **10** | -2 |
| **FES** | **1** | -13 |
| **FGFR1** | **10** | -2 |
| **FGFR1** | **1** | -8 |
| **FGFR2** | **10** | 3 |
| **FGFR2** | **1** | -4 |
| **FGFR3** | **10** | 3 |
| **FGFR3** | **1** | -3 |
| **FGFR4** | **10** | -10 |
| **FGFR4** | **1** | -5 |
| **FGR** | **10** | 1 |
| **FGR** | **1** | -7 |
| **FLT-1** | **10** | 2 |
| **FLT-1** | **1** | -4 |
| **FLT-3** | **10** | 11 |

| | | |
|---|---|---|
| **FLT-3** | **1** | -3 |
| **FLT-4** | **10** | 0 |
| **FLT-4** | **1** | -11 |
| **FMS** | **10** | 8 |
| **FMS** | **1** | -3 |
| **FYN** | **10** | 6 |
| **FYN** | **1** | -3 |
| **GAK** | **10** | 92 |
| **GAK** | **1** | 51 |
| **GLK** | **10** | -4 |
| **GLK** | **1** | -15 |
| **GRK3** | **10** | 7 |
| **GRK3** | **1** | 25 |
| **GRK5** | **10** | -8 |
| **GRK5** | **1** | -23 |
| **GRK6** | **10** | 1 |
| **GRK6** | **1** | 7 |
| **GRK7** | **10** | 1 |
| **GRK7** | **1** | -1 |
| **GSK-3-ALPHA** | **10** | -4 |
| **GSK-3-ALPHA** | **1** | -9 |
| **GSK-3-BETA** | **10** | -7 |
| **GSK-3-BETA** | **1** | -12 |
| **HASPIN** | **10** | -5 |
| **HASPIN** | **1** | -8 |
| **HCK** | **10** | -7 |
| **HCK** | **1** | -11 |
| **HIPK1** | **10** | 1 |
| **HIPK1** | **1** | -7 |
| **HIPK2** | **10** | -5 |
| **HIPK2** | **1** | -8 |
| **HIPK3** | **10** | -8 |
| **HIPK3** | **1** | -10 |
| **HIPK4** | **10** | 14 |
| **HIPK4** | **1** | 1 |
| **ICK** | **10** | 7 |
| **ICK** | **1** | -5 |
| **IGF1R** | **10** | 35 |
| **IGF1R** | **1** | -5 |
| **IKK-ALPHA** | **10** | -6 |
| **IKK-ALPHA** | **1** | -17 |
| **IKK-BETA** | **10** | -7 |
| **IKK-BETA** | **1** | 1 |
| **IKK-EPSILON** | **10** | -11 |
| **IKK-EPSILON** | **1** | -15 |
| **INSR** | **10** | -3 |

| | | |
|---|---|---|
| **INSR** | **1** | -11 |
| **IRAK1** | **10** | -11 |
| **IRAK1** | **1** | -17 |
| **IRAK4** | **10** | 2 |
| **IRAK4** | **1** | -1 |
| **IRE1** | **10** | -6 |
| **IRE1** | **1** | -17 |
| **IRR** | **10** | 39 |
| **IRR** | **1** | 5 |
| **ITK** | **10** | -1 |
| **ITK** | **1** | -4 |
| **JAK1** | **10** | -23 |
| **JAK1** | **1** | -37 |
| **JAK2** | **10** | -7 |
| **JAK2** | **1** | -11 |
| **JAK3** | **10** | 0 |
| **JAK3** | **1** | -2 |
| **JNK1** | **10** | -6 |
| **JNK1** | **1** | -8 |
| **JNK2** | **10** | -4 |
| **JNK2** | **1** | -3 |
| **JNK3** | **10** | 1 |
| **JNK3** | **1** | -4 |
| **KDR** | **10** | -26 |
| **KDR** | **1** | -22 |
| **KIT** | **10** | 14 |
| **KIT** | **1** | -2 |
| **LATS1** | **10** | 4 |
| **LATS1** | **1** | 3 |
| **LATS2** | **10** | 5 |
| **LATS2** | **1** | 2 |
| **LCK** | **10** | 3 |
| **LCK** | **1** | -3 |
| **LIMK1** | **10** | -11 |
| **LIMK1** | **1** | -17 |
| **LKB1** | **10** | -5 |
| **LKB1** | **1** | -5 |
| **LOK** | **10** | 0 |
| **LOK** | **1** | -12 |
| **LRRK2-G2019S** | **10** | 30 |
| **LRRK2-G2019S** | **1** | 9 |
| **LTK** | **10** | 4 |
| **LTK** | **1** | -2 |
| **LYNA** | **10** | 6 |
| **LYNA** | **1** | -4 |
| **LYNB** | **10** | 8 |

| | | |
|---|---|---|
| **LYNB** | **1** | -1 |
| **MAP2K4** | **10** | -4 |
| **MAP2K4** | **1** | -6 |
| **MAP2K7** | **10** | 3 |
| **MAP2K7** | **1** | -11 |
| **MAP3K8** | **10** | 48 |
| **MAP3K8** | **1** | 7 |
| **MAP4K1** | **10** | -5 |
| **MAP4K1** | **1** | -13 |
| **MAP4K2** | **10** | 10 |
| **MAP4K2** | **1** | -3 |
| **MAP4K3** | **10** | 18 |
| **MAP4K3** | **1** | 16 |
| **MAP4K4** | **10** | 15 |
| **MAP4K4** | **1** | -2 |
| **MAP4K5** | **10** | 11 |
| **MAP4K5** | **1** | -4 |
| **MAPK1** | **10** | 4 |
| **MAPK1** | **1** | -4 |
| **MAPK3** | **10** | 4 |
| **MAPK3** | **1** | -3 |
| **MAPKAPK-2** | **10** | 1 |
| **MAPKAPK-2** | **1** | -4 |
| **MAPKAPK-3** | **10** | -2 |
| **MAPKAPK-3** | **1** | -12 |
| **MARK1** | **10** | 23 |
| **MARK1** | **1** | -8 |
| **MARK3** | **10** | 13 |
| **MARK3** | **1** | 5 |
| **MARK4** | **10** | 3 |
| **MARK4** | **1** | 2 |
| **MEK1** | **10** | 3 |
| **MEK1** | **1** | -3 |
| **MEK2** | **10** | 3 |
| **MEK2** | **1** | -2 |
| **MEK3** | **10** | 4 |
| **MEK3** | **1** | -1 |
| **MEKK3** | **10** | -21 |
| **MEKK3** | **1** | -41 |
| **MELK** | **10** | 6 |
| **MELK** | **1** | -2 |
| **MER** | **10** | 27 |
| **MER** | **1** | 2 |
| **MET** | **10** | -1 |
| **MET** | **1** | -4 |
| **MINK** | **10** | -13 |

| | | |
|---|---|---|
| **MINK** | **1** | -21 |
| **MLK1** | **10** | -10 |
| **MLK1** | **1** | -21 |
| **MLK2** | **10** | -24 |
| **MLK2** | **1** | -35 |
| **MLK3** | **10** | 5 |
| **MLK3** | **1** | 0 |
| **MNK1** | **10** | -7 |
| **MNK1** | **1** | -11 |
| **MNK2** | **10** | -9 |
| **MNK2** | **1** | -17 |
| **MRCKA** | **10** | 6 |
| **MRCKA** | **1** | 2 |
| **MRCKB** | **10** | 0 |
| **MRCKB** | **1** | -4 |
| **MSK1** | **10** | 2 |
| **MSK1** | **1** | -6 |
| **MSK2** | **10** | 0 |
| **MSK2** | **1** | -3 |
| **MSSK1** | **10** | 12 |
| **MSSK1** | **1** | -2 |
| **MST1** | **10** | 7 |
| **MST1** | **1** | -2 |
| **MST2** | **10** | 3 |
| **MST2** | **1** | -4 |
| **MST3** | **10** | 17 |
| **MST3** | **1** | -1 |
| **MST4** | **10** | 34 |
| **MST4** | **1** | 0 |
| **mTOR** | **10** | 0 |
| **mTOR** | **1** | -2 |
| **mTORC1** | **10** | 1 |
| **mTORC1** | **1** | 0 |
| **MUSK** | **10** | 5 |
| **MUSK** | **1** | -1 |
| **MYO3A** | **10** | 5 |
| **MYO3A** | **1** | 8 |
| **MYO3B** | **10** | -2 |
| **MYO3B** | **1** | 2 |
| **NDR1** | **10** | 3 |
| **NDR1** | **1** | -1 |
| **NDR2** | **10** | 3 |
| **NDR2** | **1** | 0 |
| **NEK1** | **10** | 9 |
| **NEK1** | **1** | -20 |
| **NEK2** | **10** | -5 |

| | | |
|---|---|---|
| **NEK2** | **1** | -11 |
| **NEK3** | **10** | 5 |
| **NEK3** | **1** | 5 |
| **NEK4** | **10** | 1 |
| **NEK4** | **1** | -18 |
| **NEK5** | **10** | -2 |
| **NEK5** | **1** | -7 |
| **NEK6** | **10** | -3 |
| **NEK6** | **1** | -8 |
| **NEK7** | **10** | -2 |
| **NEK7** | **1** | -5 |
| **NEK9** | **10** | -8 |
| **NEK9** | **1** | -9 |
| **NIM1K** | **10** | -4 |
| **NIM1K** | **1** | 0 |
| **NUAK2** | **10** | 2 |
| **NUAK2** | **1** | -7 |
| **P38-ALPHA** | **10** | -2 |
| **P38-ALPHA** | **1** | -7 |
| **P38-BETA** | **10** | 1 |
| **P38-BETA** | **1** | -9 |
| **P38-DELTA** | **10** | -6 |
| **P38-DELTA** | **1** | -9 |
| **P38-GAMMA** | **10** | 0 |
| **P38-GAMMA** | **1** | -4 |
| **P70S6K1** | **10** | -9 |
| **P70S6K1** | **1** | -13 |
| **P70S6K2** | **10** | 16 |
| **P70S6K2** | **1** | -6 |
| **PAK1** | **10** | 4 |
| **PAK1** | **1** | -2 |
| **PAK2** | **10** | -1 |
| **PAK2** | **1** | -5 |
| **PAK3** | **10** | -3 |
| **PAK3** | **1** | -2 |
| **PAK4** | **10** | 0 |
| **PAK4** | **1** | -1 |
| **PAK5** | **10** | 0 |
| **PAK5** | **1** | -1 |
| **PAK6** | **10** | 0 |
| **PAK6** | **1** | -1 |
| **PAR1BA** | **10** | 5 |
| **PAR1BA** | **1** | 2 |
| **PASK** | **10** | 0 |
| **PASK** | **1** | -7 |
| **PBK** | **10** | -2 |

| | | |
|---|---|---|
| **PBK** | **1** | -15 |
| **PDGFR-ALPHA** | **10** | 19 |
| **PDGFR-ALPHA** | **1** | 4 |
| **PDGFR-BETA** | **10** | 13 |
| **PDGFR-BETA** | **1** | 1 |
| **PDK1** | **10** | -5 |
| **PDK1** | **1** | -7 |
| **PDK2** | **10** | -17 |
| **PDK2** | **1** | -31 |
| **PDK3** | **10** | 29 |
| **PDK3** | **1** | 0 |
| **PDK4** | **10** | 16 |
| **PDK4** | **1** | 4 |
| **PHK-GAMMA1** | **10** | -2 |
| **PHK-GAMMA1** | **1** | -7 |
| **PHK-GAMMA2** | **10** | 0 |
| **PHK-GAMMA2** | **1** | -6 |
| **PI3K-ALPHA** | **10** | 23 |
| **PI3K-ALPHA** | **1** | 46 |
| **PI3K-BETA** | **10** | 0 |
| **PI3K-BETA** | **1** | 10 |
| **PI3K-DELTA** | **10** | 24 |
| **PI3K-DELTA** | **1** | 11 |
| **PI3K-GAMMA** | **10** | -2 |
| **PI3K-GAMMA** | **1** | -3 |
| **PI4-K-BETA** | **10** | -7 |
| **PI4-K-BETA** | **1** | -8 |
| **PIKFYVE** | **10** | 55 |
| **PIKFYVE** | **1** | 17 |
| **PIM-1-KINASE** | **10** | 1 |
| **PIM-1-KINASE** | **1** | -2 |
| **PIM2** | **10** | -1 |
| **PIM2** | **1** | -5 |
| **PIM3** | **10** | 3 |
| **PIM3** | **1** | 7 |
| **PKA** | **10** | 1 |
| **PKA** | **1** | 1 |
| **PKAC-BETA** | **10** | 1 |
| **PKAC-BETA** | **1** | -2 |
| **PKAC-GAMMA** | **10** | -3 |
| **PKAC-GAMMA** | **1** | -3 |
| **PKC-ALPHA** | **10** | 16 |
| **PKC-ALPHA** | **1** | 2 |
| **PKC-BETA1** | **10** | 2 |
| **PKC-BETA1** | **1** | 1 |
| **PKC-BETA2** | **10** | -9 |

| | | |
|---|---|---|
| **PKC-BETA2** | **1** | -8 |
| **PKC-DELTA** | **10** | -4 |
| **PKC-DELTA** | **1** | -11 |
| **PKC-EPSILON** | **10** | 1 |
| **PKC-EPSILON** | **1** | -4 |
| **PKC-ETA** | **10** | -12 |
| **PKC-ETA** | **1** | -4 |
| **PKC-GAMMA** | **10** | -6 |
| **PKC-GAMMA** | **1** | -9 |
| **PKC-IOTA** | **10** | -7 |
| **PKC-IOTA** | **1** | -5 |
| **PKC-THETA** | **10** | -1 |
| **PKC-THETA** | **1** | -3 |
| **PKC-ZETA** | **10** | 5 |
| **PKC-ZETA** | **1** | -1 |
| **PKN1** | **10** | -23 |
| **PKN1** | **1** | -4 |
| **PKN2** | **10** | -15 |
| **PKN2** | **1** | 2 |
| **PKN3** | **10** | 5 |
| **PKN3** | **1** | -2 |
| **PLK1** | **10** | 0 |
| **PLK1** | **1** | -10 |
| **PLK2** | **10** | 4 |
| **PLK2** | **1** | 2 |
| **PLK3** | **10** | -2 |
| **PLK3** | **1** | -3 |
| **PLK4** | **10** | 9 |
| **PLK4** | **1** | -10 |
| **PRAK** | **10** | 2 |
| **PRAK** | **1** | -5 |
| **PRKD1** | **10** | 17 |
| **PRKD1** | **1** | -10 |
| **PRKD2** | **10** | 27 |
| **PRKD2** | **1** | -9 |
| **PRKD3** | **10** | 8 |
| **PRKD3** | **1** | -8 |
| **PRKG1** | **10** | 3 |
| **PRKG1** | **1** | -3 |
| **PRKG2** | **10** | 8 |
| **PRKG2** | **1** | 0 |
| **PRKX** | **10** | 0 |
| **PRKX** | **1** | -1 |
| **PTK5** | **10** | 6 |
| **PTK5** | **1** | -1 |
| **PYK2** | **10** | -2 |

| | | |
|---|---|---|
| **PYK2** | **1** | -6 |
| **QIK** | **10** | -6 |
| **QIK** | **1** | -11 |
| **RET** | **10** | 10 |
| **RET** | **1** | -1 |
| **RIPK1** | **10** | -11 |
| **RIPK1** | **1** | -49 |
| **RIPK2** | **10** | 3 |
| **RIPK2** | **1** | -3 |
| **ROCK1** | **10** | -1 |
| **ROCK1** | **1** | -1 |
| **ROCK2** | **10** | -1 |
| **ROCK2** | **1** | -3 |
| **RON** | **10** | 15 |
| **RON** | **1** | 3 |
| **ROS** | **10** | 3 |
| **ROS** | **1** | -3 |
| **RSK1** | **10** | 2 |
| **RSK1** | **1** | -1 |
| **RSK2** | **10** | 1 |
| **RSK2** | **1** | -2 |
| **RSK3** | **10** | 9 |
| **RSK3** | **1** | -7 |
| **RSK4** | **10** | -8 |
| **RSK4** | **1** | -11 |
| **SGK1** | **10** | -1 |
| **SGK1** | **1** | -3 |
| **SGK2** | **10** | 9 |
| **SGK2** | **1** | -1 |
| **SGK3** | **10** | 0 |
| **SGK3** | **1** | 0 |
| **SIK** | **10** | 0 |
| **SIK** | **1** | -5 |
| **SLK** | **10** | 21 |
| **SLK** | **1** | -5 |
| **SPHK1** | **10** | 32 |
| **SPHK1** | **1** | 27 |
| **SPHK2** | **10** | 6 |
| **SPHK2** | **1** | 11 |
| **SRC** | **10** | -3 |
| **SRC** | **1** | -16 |
| **SRMS** | **10** | -1 |
| **SRMS** | **1** | -2 |
| **SRPK1** | **10** | 3 |
| **SRPK1** | **1** | -5 |
| **SRPK2** | **10** | 22 |

| | | |
|---|---|---|
| **SRPK2** | **1** | -8 |
| **SSTK** | **10** | 2 |
| **SSTK** | **1** | -9 |
| **STK16** | **10** | 8 |
| **STK16** | **1** | 1 |
| **STK25** | **10** | 33 |
| **STK25** | **1** | 13 |
| **STK33** | **10** | 4 |
| **STK33** | **1** | -2 |
| **SYK** | **10** | 0 |
| **SYK** | **1** | -7 |
| **TAK1-TAB1** | **10** | 12 |
| **TAK1-TAB1** | **1** | 8 |
| **TAOK2** | **10** | 0 |
| **TAOK2** | **1** | -8 |
| **TAOK3** | **10** | -15 |
| **TAOK3** | **1** | -20 |
| **TBK1** | **10** | -9 |
| **TBK1** | **1** | -12 |
| **TEC** | **10** | -1 |
| **TEC** | **1** | -4 |
| **TESK1** | **10** | -4 |
| **TESK1** | **1** | 0 |
| **TIE2** | **10** | 5 |
| **TIE2** | **1** | 0 |
| **TLK1** | **10** | -2 |
| **TLK1** | **1** | -5 |
| **TNIK** | **10** | 10 |
| **TNIK** | **1** | -12 |
| **TNK1** | **10** | 5 |
| **TNK1** | **1** | 2 |
| **TNK2** | **10** | 0 |
| **TNK2** | **1** | 0 |
| **TRKA** | **10** | 4 |
| **TRKA** | **1** | -2 |
| **TRKB** | **10** | 5 |
| **TRKB** | **1** | 0 |
| **TRKC** | **10** | 9 |
| **TRKC** | **1** | -2 |
| **TSSK1** | **10** | 0 |
| **TSSK1** | **1** | -5 |
| **TSSK2** | **10** | -2 |
| **TSSK2** | **1** | -6 |
| **TSSK3** | **10** | -3 |
| **TSSK3** | **1** | -4 |
| **TTBK1** | **10** | 4 |

| | | |
|---|---|---|
| **TTBK1** | **1** | -7 |
| **TTBK2** | **10** | 3 |
| **TTBK2** | **1** | -1 |
| **TTK** | **10** | -3 |
| **TTK** | **1** | 0 |
| **TXK** | **10** | -2 |
| **TXK** | **1** | -9 |
| **TYK2** | **10** | 3 |
| **TYK2** | **1** | 2 |
| **TYRO3** | **10** | 5 |
| **TYRO3** | **1** | -3 |
| **ULK1** | **10** | 2 |
| **ULK1** | **1** | -7 |
| **ULK3** | **10** | -3 |
| **ULK3** | **1** | -3 |
| **WEE1** | **10** | 13 |
| **WEE1** | **1** | -9 |
| **WNK1** | **10** | -14 |
| **WNK1** | **1** | -23 |
| **YES** | **10** | 7 |
| **YES** | **1** | -4 |
| **ZAP70** | **10** | 36 |
| **ZAP70** | **1** | 10 |
